# Monocytes Mobilized by Gut Neurons Remodel the Enteric Nervous System

**DOI:** 10.1101/2024.06.09.598124

**Authors:** Sravya Kurapati, Changsik Shin, Krisztina Szabo, Yu Liu, Azree Zaffran Ashraf, Balazs Koscso, Chinmayee Dash, Leonardo Navarro, Monalee Saha, Sushma Nagaraj, Wenhui Wang, Jun Zhu, Natalya Shulzhenko, Christina E. Baer, Shanthi Srinivasan, Subhash Kulkarni, Pankaj Jay Pasricha, Lauren A. Peters, Milena Bogunovic

**Affiliations:** Department of Pathology, University of Massachusetts Chan Medical School, Worcester, MA; The Biomedical Sciences PhD Program, Penn State University College of Medicine, Hershey, PA; Department of Microbiology and Immunology, Penn State University College of Medicine, Hershey, PA; Department of Genetics & Genomic Sciences, Icahn School of Medicine at Mount Sinai, New York, NY; The Morningside Graduate School of Biomedical Sciences at UMass Chan Medical School, Worcester, MA; Center for Neurogastroenterology, The Johns Hopkins University School of Medicine, Baltimore, MA; College of Veterinary Medicine, Oregon State University, Corvallis, OR; Sanderson Center for Optical Experimentation (SCOPE), Department of Microbiology and Physiological Systems, University of Massachusetts Chan Medical School, Worcester, MA; Division of Digestive Diseases, Department of Medicine, Emory University School of Medicine, Atlanta, GA; Division of Gastroenterology, Department of Medicine, Beth Israel Deaconess Medical Center, Harvard Medical School, Boston, MA; Department of Medicine, Mayo Clinic, Phoenix, AZ; Windreich Department of AI & Human Health, Icahn School of Medicine at Mount Sinai

## Abstract

The proper organization of the enteric nervous system (ENS) is critical for normal gastrointestinal (GI) physiology. Inflammatory bowel disease (IBD) dysregulates GI physiology, including bowel movements (motility), but in many IBD patients, GI motility disorders persist in remission through a poorly understood pathological process. Here we uncover that post-inflammatory GI dysmotility (PI-GID) stems from structural ENS remodeling driven by a combination of neuronal loss and neurogenesis. Enteric neurons respond to mucosal inflammation by upregulating CCL2 expression and facilitating the recruitment of CCR2^+^ monocytes into the neural myenteric plexus within the intestinal muscle. This is followed by the expansion of monocyte-derived macrophages and their migration into the myenteric ganglia and phagocytosis of neurons. However, excessive recruitment of monocytes results in disproportionate ENS remodeling and PI-GID. The expansion of inflammatory cells is known to promote tissue hypoxia. We find that enteric neurons become hypoxic upon colitis, but hypoxia-induced signaling via HIF1α initiates an adaptation program in enteric neurons to attenuate CCL2 expression and limit monocyte recruitment. We demonstrate that reinforcing HIF1α signaling in enteric neurons prevents PI-GID by reducing colitis-associated monocyte recruitment in the myenteric plexus and protecting against ENS remodeling. In summary, our findings unveil PI-GID pathogenesis and identify a regulatory axis for its prevention.

**One Sentence Summary:** Intestinal mucosal inflammation engages enteric neurons in the inflammatory response leading to neurogenic recruitment of monocytes into the extra-mucosal myenteric plexus followed by pathological structural remodeling of the enteric nervous system by monocyte-derived macrophages.

## INTRODUCTION

Inflammatory bowel disease (IBD) which includes Crohn’s disease (CD) and ulcerative colitis (UC) is a progressive inflammatory condition of the gastrointestinal (GI) tract of non-infectious etiology. Great strides have been made in the treatment of active inflammation in patients with IBD. However, many patients with IBD in remission develop functional GI disorders including irritable bowel syndrome (IBS), and present with a wide range of symptoms such as chronic abdominal pain and various GI motility dysfunctions (dysmotility) afflicting both upper and lower gut regions^1–5^. And yet, there is a lack of understanding of how IBD permanently dysregulates key GI functions.

The GI tract performs vital physiological functions related to food and water consumption. Its dense innervation by the central and peripheral nervous systems allows it to carry out complex physiological processes in a highly coordinated manner. Most gut-innervating peripheral neurons are located within the gut tissue, where they form the enteric nervous system (ENS), the second largest collection of neurons outside of the brain^6^. The vital physiological GI functions (smooth muscle contractility, epithelial barrier integrity, and blood flow) and some aspects of its mucosal immune response are tightly regulated by the ENS^6^. Of the several layers of the gut, the intestinal mucosa is the most proximal to the lumen and is exposed to various stressors, while the somas of enteric neurons are positioned outside of the mucosa in two major plexuses, submucosal and myenteric, with preferential control of the epithelial barrier and smooth muscle contractions, respectively^7^. Significant advances have been made in our understanding of mucosal immune responses in IBD, but extra-mucosal cellular GI compartments, specifically the ENS, have received very little attention. So, it remains largely unknown how mucosal inflammation affects ENS homeostasis and function in IBD^1^.

A unique population of intestinal macrophages named muscularis macrophages (MMs) resides in the outer smooth muscle layer of the intestine (muscularis externa) where it anatomically associates with nerve fibers and somas of myenteric neurons^8–10^. Macrophages are immune cells with “housekeeping” functions^11^, and accumulating evidence points to a symbiotic relationship between MMs and enteric neurons in early postnatal ENS development and homeostasis of adult ENS in health and GI infection^12^, but the functional connection between the two cell types in IBD is yet to be identified.

Here we sought to establish the mechanisms of post-inflammatory GI dysmotility (PI-GID). By testing the hypothesis that mucosal inflammation in IBD alters homeostatic crosstalk between enteric neurons and MMs, we uncovered a colitis-driven neuroimmune pathway underlying pathological remodeling of the ENS with associated dysfunction and identified its regulatory axis.

## RESULTS

### Transient colitis results in long-term post-inflammatory GI dysmotility

IBD is a chronic inflammatory condition with a relapsing-remitting pattern. To study PI-GID, we established a model of transient colitis to reflect phases of active mucosal inflammation and post-inflammatory remission based on the chronic DSS colitis model^13^. Eight-week-old C57BL/6 wild-type (WT) mice were given three 5-day cycles of DSS to maintain active colitis for four weeks (3×DSS colitis); then mice were followed for at least another six to eight weeks until a nearly complete recovery to reproduce remission in patients with IBD (Fig. 1a). The DSS phase was associated with significant body weight loss, increased fecal lipocalin-2 concentration as a biomarker of mucosal inflammation^14^, and increased mucosal macrophage and neutrophil counts, consistent with active mucosal inflammation. At the late post-inflammatory phase, there was no difference in body weight, fecal lipocalin-2 levels and mucosal macrophage counts between the DSS and water groups although the mucosal neutrophil counts remained slightly increased (Fig. 1b-e). These changes overall characterize the recovery phase. In the follow up experiments, mucosal neutrophil counts were used as the most sensitive readout of mucosal inflammation. Despite a nearly complete recovery from colitis, mice with a history of DSS treatment had signs of GI dysmotility as evidenced by accelerated GI transit (Fig. 1f). Increased tone of intestinal smooth muscle at the post-inflammatory phase was reflected in the shortened colon, despite a significant recovery as compared to the DSS phase (Fig. 1g). Therefore, functional changes in the 3×DSS colitis model overlap with the changes observed in patients with PI-GID^1^.

**Figure 1.**
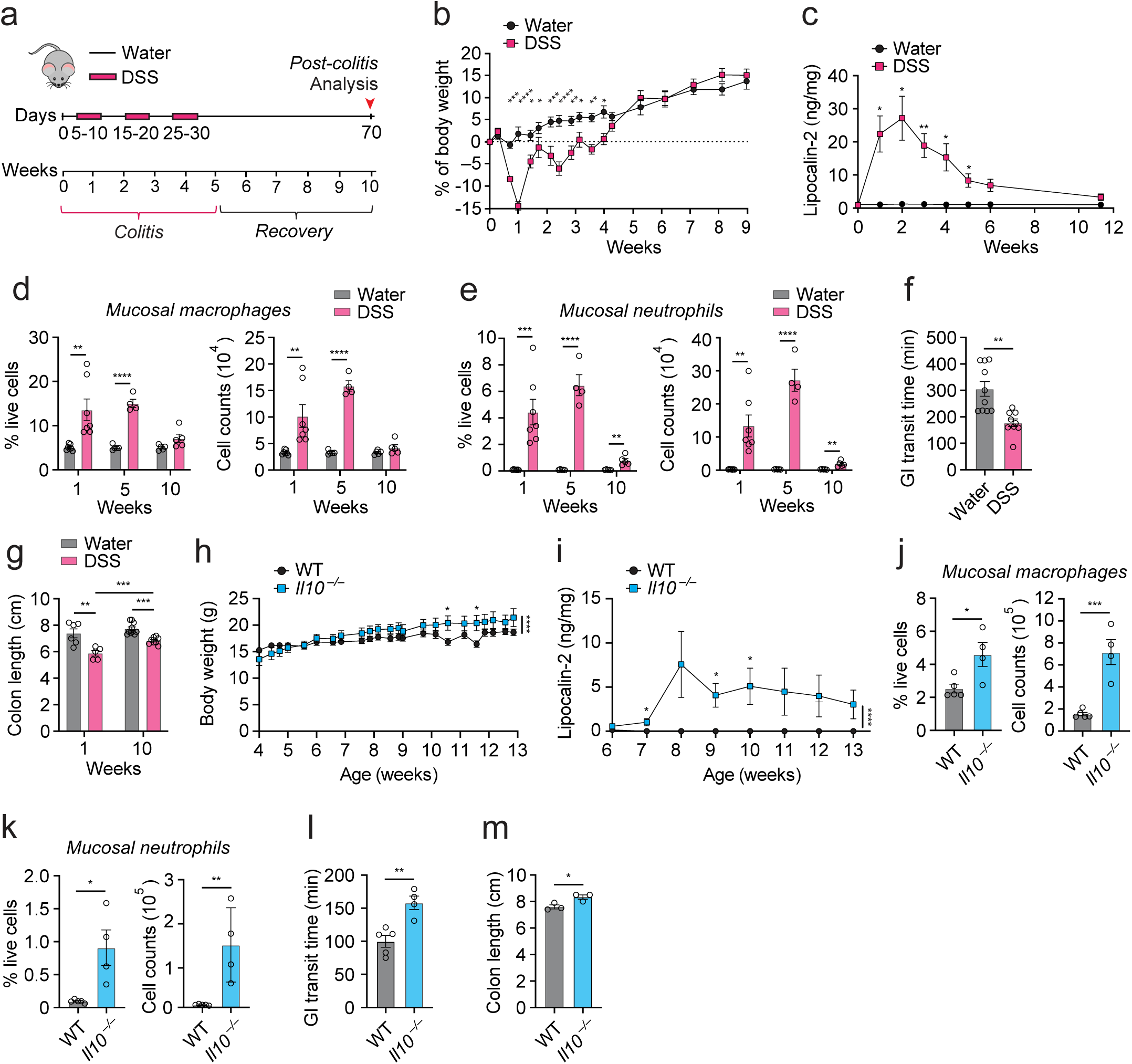
Transient colitis leads to long-term post-inflammatory gastrointestinal dysmotility (PI-GID). (a) Experimental design of transient 3×DSS colitis. (b) Percentage of body weight change in 3×DSS- or water-treated WT mice. (c) Fecal lipocalin-2 in 3×DSS- or water-treated WT mice, measured by ELISA. (d) Percentage of total viable cells and absolute cell counts of colonic mucosal macrophages in 3×DSS- or water-treated mice, quantified by flow cytometry (FC). (e) Percentage of total viable cells and absolute cell counts of colonic mucosal neutrophils in 3×DSS- or water-treated mice, quantified by FC. (f) Gastrointestinal (GI) transit in 3×DSS- or water-treated mice at week 12 after starting DSS. (g) Colon length in 3×DSS- or water-treated mice. (h) Body weight in *Il10*^−/−^ mice from 4 weeks of age until they developed rectal prolapse or in WT littermates. (i) Fecal lipocalin-2 in *Il10*^−/−^ and WT mice, measured by ELISA. (j) Percentage of total viable cells and absolute cell counts of colonic mucosal macrophages in 13-week-old *Il10*^−/−^ and WT mice, quantified by FC. (k) Percentage of total viable cells and absolute cell counts of colonic mucosal neutrophils in 13-week-old *Il10*^−/−^ and WT mice, quantified by FC. (l) GI transit in 13-week-old *Il10*^−/−^ and WT mice. (m) Colon length in 13-week-old *Il10*^−/−^ and WT mice. All graphs show mean ± SEM from 1 representative experiment. Statistical analyses: Unpaired Student’s t-test and two-way ANOVA for multiple comparisons, *P≤0.05, **P≤0.01, ***P≤0.001, and ****P≤0.0001.

As opposed to mice with transient colitis, age-matched *Il10*^−/−^ mice susceptible to spontaneous colitis^15^ had steadily increased fecal lipocalin-2 levels, and significantly higher mucosal macrophage and neutrophil counts at the late time point (Fig. 1h-k) indicative of progressive inflammation. *Il10*^−/−^ mice also had signs of GI dysmotility, although in contrast to the 3×DSS colitis, the GI transit in *Il10*^−/−^ mice was delayed (Fig. 1l) suggesting different underlying mechanisms. The elongated colon (Fig. 1m) along with signs of rectal prolapse indicated reduced smooth muscle tone. The presence of dysmotility in both models pointed to functionally altered ENS. Therefore, we focused on elucidating the changes in the ENS caused by colitis.

### Transient colitis induces structural remodeling of the ENS through a combination of neuronal loss and neurogenesis

To understand the nature of ENS changes caused by colitis we imaged the colonic myenteric plexus at different stages of colitis. Littermate mice were treated with 3×DSS or water (control). The colonic muscularis externa was isolated at the early acute (weeks 1 and 3) and late post-inflammatory (after week 10) time points, and the same regions of the colonic myenteric plexus were stained with the antibodies against Hu (B, C and D subunits)^16^, ýIII-Tubulin and MHC Class II (MHCII) to visualize neuronal somas, nerve fiber network, and macrophages, respectively, and imaged by confocal microscopy (Fig. 2a). The total number of myenteric neurons, neuronal density, and nerve fiber architecture were compared between the groups. We found significant structural changes in the myenteric plexus of mice with colitis, and the pattern of these changes differed between the early and late time points as colitis progressed and regressed (Fig. 2b). At week 1, colitis was associated with a significant reduction in the total number of myenteric neurons, the breakdown of larger ganglia into smaller clusters, and structural changes in the nerve fiber network caused by the partial loss of inter-ganglionic nerve fiber tracts (IGFTs) (see Methods for quantification approach). These changes were more evident at week 3 of colitis. In contrast, at the late post-inflammatory phase, the total number of myenteric neurons recovered in the 3×DSS group, although there was a large variation in neuronal counts among the fields with the areas of severe neuronal loss alternated by hyperplastic ganglia. Further, the density of the IGFT network was increased. The changes in the myenteric plexus architecture could not be explained by shifts in the organ surface area because the colon remained shortened across all time points (Fig. 1g).

**Figure 2.**
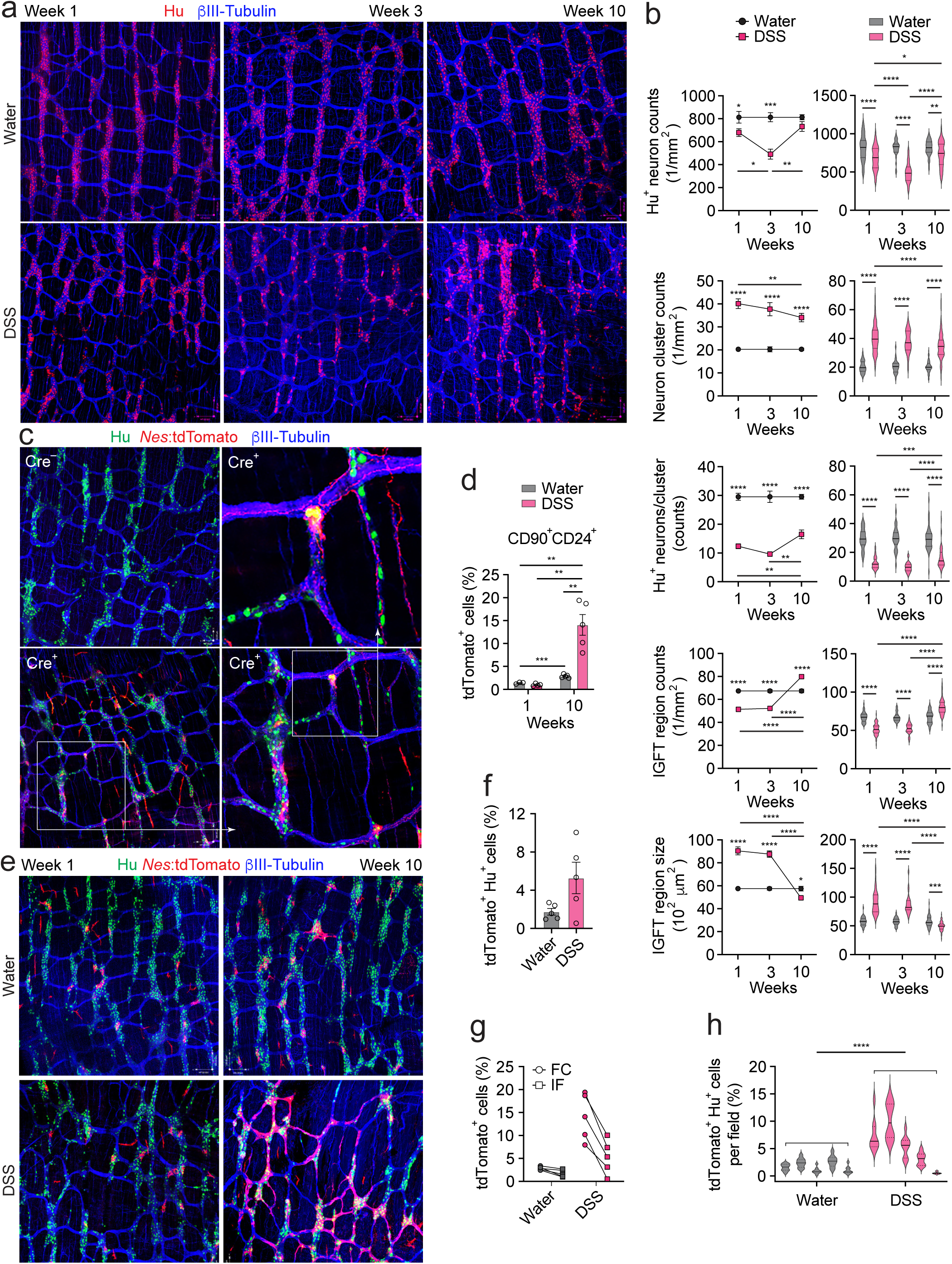
Transient colitis induces neuronal loss and neurogenesis that facilitate structural remodeling of the myenteric plexus. (a) Confocal images of colonic myenteric plexus isolated from 3×DSS- or water-treated mice and stained with Hu (neuronal somas), ýIII-Tubulin (nerve fibers) and MHCII (MMs, channel turned off) antibodies. Scale bars, 100 μm. (b) Structural changes in myenteric plexus from mice described in (a) quantified by confocal microscopy as explained in Methods. Data shown as mean value per mouse per group (left) or as distribution of pooled values per group (right). (c) Confocal images of colonic myenteric plexus isolated from tamoxifen- and 1×DSS- or water-treated *Nes*^ER-Cre^*Rosa26*(*R26*)-STOP^fl/fl^tdTomato (Cre*^+^*) and control *R26*-STOP^fl/fl^tdTomato (Cre^−^) mice on day 45 (6.5 weeks), stained with antibodies to Hu (neuronal somas), tdTomato (newborn cells) and βIII-Tubulin (nerve fibers). Scale bars, 62 μm. (d) Percentage of newly generated tdTomato^+^ cells among CD90^+^CD24^+^ colonic myenteric neurons isolated from tamoxifen- and 3×DSS- or water-treated *Nes*^ER-Cre^*R26*-STOP^fl/fl^tdTomato (Cre^+^) mice, quantified by FC. (e) Confocal images of colonic myenteric plexus from Cre^+^ mice described in (d). Scale bars, 100 μm. (f) Percentage of tdTomato^+^Hu ^+^ neurons in the colonic myenteric plexus from Cre*^+^* mice described in (d) at week 10, quantified by confocal microscopy. (g) Pair-wise comparison of percentages of newly generated tdTomato^+^ myenteric neurons quantified by FC and confocal microscopy in Cre*^+^*mice described in (d) at week 10. (h) Percentage distribution of newly generated tdTomato^+^Hu^+^ colonic myenteric neurons per field per mouse in Cre*^+^*mice described in (d) at week 10, quantified by confocal microscopy. Each violin plot represents a mouse. All graphs show mean ± SEM from 1 representative experiment. Statistical analyses: Unpaired Student’s t-test and two-way ANOVA for multiple comparisons, *P≤0.05, **P≤0.01, ***P≤0.001, and ****P≤0.0001.

A partial loss of enteric neurons due to programmed cell death has been described in the acute phase of colitis^12,17^, and a long-term study in infectious colitis showed progressive neuronal loss^12^. We therefore used a second approach to confirm our results. CD90- and CD24-specific antibodies have been used to stain enteric neurons in human tissue^18^. Thus, using flow cytometry we established that a single-cell suspension of murine muscularis externa contained the CD45^−^ non-hematopoietic CD90^+^CD24^+^ cell subset positive for a pan-neuronal marker PGP9.5 (Extended Data Fig.1a,b). CD90^+^CD24^+^ cells also selectively expressed neuron-specific genes *Elavl3* (encodes Hu C subunit) and *Elavl4* (encodes Hu D subunit) (Extended Data Fig. 1c). Next, using flow cytometry, we quantified the number of CD90^+^CD24^+^ cells in the muscularis externa of mice with 3×DSS colitis and controls, and found the results to be similar to confocal microscopy – the reduction of CD90^+^CD24^+^ cells in the DSS group was transient (Extended Data Fig. 1d). Therefore, neuronal loss in response to mucosal inflammation is reversed post-colitis, but it is unclear what causes the recovery of neuronal counts.

Given the observed neuronal hyperplasia in some areas of the myenteric plexus post-colitis, we hypothesized that enteric neurogenesis takes place. Nestin^+^ enteric neural precursor cells (ENPCs) have been described as progenitors of adult enteric neurons^19,20,21^ although the extent of neurogenesis in the adult intestine remains unclear^7^. To test the contribution of ENPC-driven neurogenesis to colitis-induced ENS remodeling, we used a described fate-mapping approach^19^ based on tamoxifen-treated *Nestin*(*Nes*)^ER-Cre^*Rosa26*-STOP^fl/fl^tdTomato (*Nes*:tdTomato) mice to label newborn neurons under conditions of colitis. After only one cycle of DSS, we were able to detect new Hu^+^tdTomato^+^ neurons at the post-colitis phase with morphology ranging from less differentiated star-like shaped cells to fully formed neurons with axons embedded into the myenteric plexus (Fig. 2c). Other tdTomato^+^ cell types included intra-ganglionic S100ý^+^ enteric glia cells and extra-ganglionic perivascular cells (Extended Data Fig. 1e). Next, we quantified the number of Nestin^+^ ENPC-derived neurons in the 3×DSS model. By flow cytometry, about 15% of neuron-enriched CD90^+^CD24^+^ cells were tdTomato^+^ at the late post-inflammatory phase (Fig. 2d, Extended Data Fig. 1f,g). Similar results were found by confocal microscopy (Fig. 2e-g), although with more variation among individual mice and fields (Fig. 2h), likely reflecting the non-homogeneous impact of colitis along the colon and the subsequent non-uniform ENS remodeling. Collectively, in transient colitis, increased Nestin^+^ ENPC-driven neurogenesis provides the biological basis for the recovery of neuronal counts and remodeled ENS architecture post-colitis.

In contrast to transient colitis, *Il10*^−/−^ mice with progressive colitis showed a combination of neuronal loss and increased density of the IGFT network likely reflecting remodeling due to neurogenesis (Extended Data Fig. 1h,i). This suggested that the rate of neuronal loss exceeds neurogenesis in this model, potentially explaining why GI transit time was delayed in these mice^12^. Because of our focus on PI-GID, we did not investigate the ENS changes in *Il10*^−/−^ mice further and, in the follow-up experiments, focused on the 3×DSS model.

### Monocyte-derived muscularis macrophages with microglia-like properties expand in the myenteric plexus in response to colitis

The cellular mechanisms driving colitis-induced ENS remodeling are not known. Macrophages in other organ systems have been shown to orchestrate biological processes relevant to tissue remodeling including the safe disposal of cell debris through efferocytosis and the release of regulatory mediators and growth factors^11^. We, therefore, hypothesized that ENS-associated MMs, shown to support postnatal and adult enteric neurons^8,9,12,22,23^, are the cell type responsible for colitis-induced ENS remodeling and sought to define the nature of changes in the MM compartment upon transient colitis. Quantification of MMs in the myenteric plexus by confocal microscopy and flow cytometry at the early acute and late post-inflammatory time points of 3×DSS colitis revealed a transient increase in their number (Fig. 3a-c).

**Figure 3.**
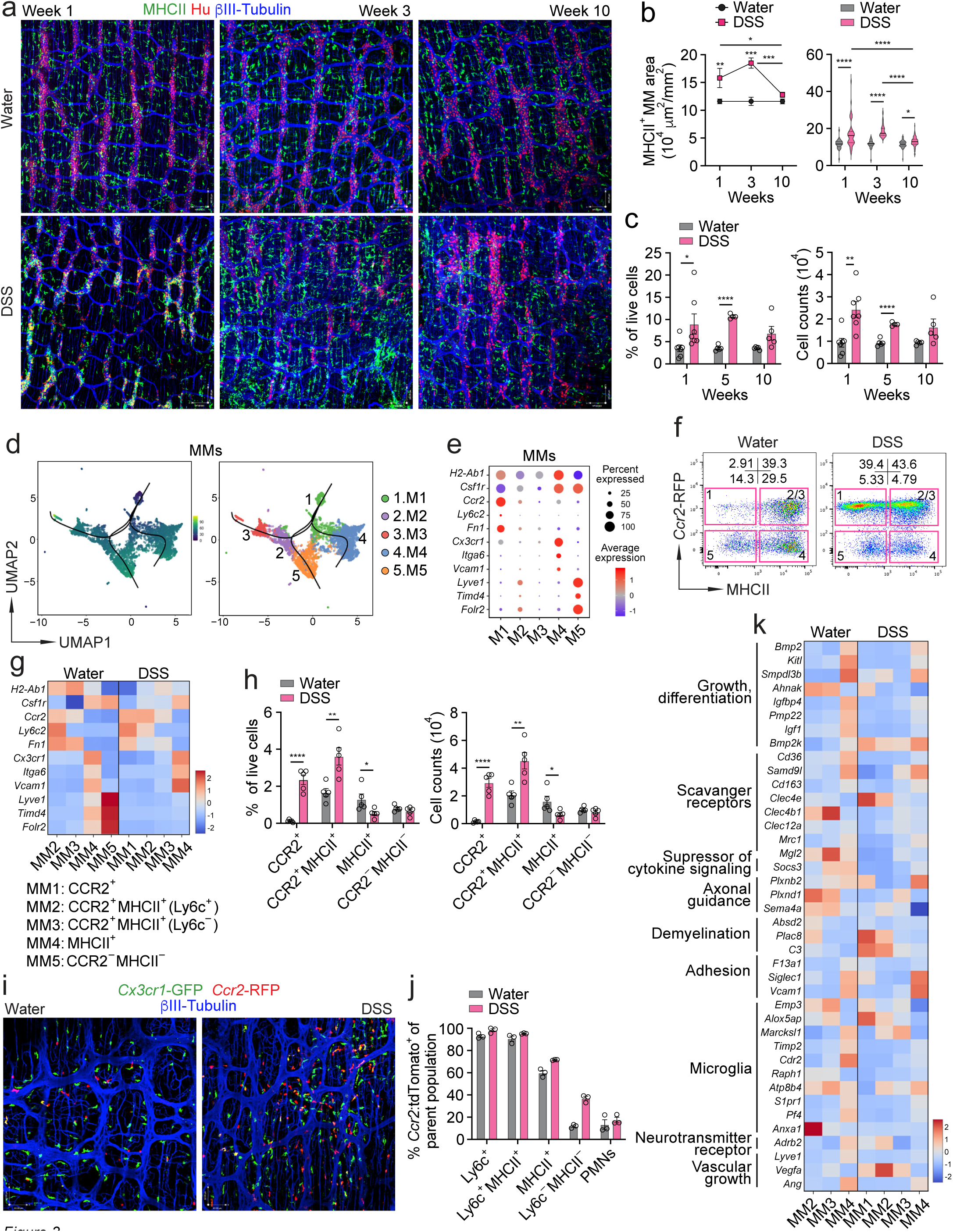
CCR2^+^ monocyte-derived muscularis macrophages (MMs) expand in the myenteric plexus in response to colitis. (a) Confocal images of colonic myenteric plexus isolated from 3×DSS- or water-treated mice (as in Fig. 2a but with MHCII channel on). Scale bars, 100 μm. (b) Total MHCII^+^ area in the colonic myenteric plexus shown in (a), quantified by confocal microscopy. Data are shown as mean value per mouse per group (left) or as a distribution of pooled values per group (right). (c) Percentage of total viable cells (left) and absolute cell counts (right) of colonic MMs from mice described in (a), quantified by FC. (d) Predicted developmental trajectory of five scRNA-seq MM clusters in normal colon determined by slingshot analysis, shown as pseudotime (left) and MM clusters (right). (e) ScRNA-seq dot heatmap showing expression of 11 marker genes across five MM clusters identified in (d). Color denotes the average expression level and size represents the percentage of cells expressing the indicated genes. (f) FC plots showing colonic MM subsets and their percentages in 1×DSS- or water-treated *Ccr2*^RFP/+^*Cx3cr1*^GFP/+^ reporter mice at week 1. Gated on viable CD45^+^CD11b^+^CD16/32^+^ cells. (g) Bulk RNA-seq heatmap showing expression of 11 marker genes by MM subsets defined in (f) isolated by four-way FACS from 1×DSS- or water-treated *Ccr2*^RFP/+^*Cx3cr1*^GFP/+^ mice at week 1. (h) Percentage of total viable cells (left) and absolute counts (right) of colonic MM subsets shown in (f), quantified by FC. (i) Confocal images of colonic myenteric plexus from 1×DSS- or water-treated *Ccr2*^RFP/+^*Cx3cr1*^GFP/+^ mice at week 1. Scale bars, 62 μm. (j) Percentage of monocyte-derived tdTomato^+^ cells among colonic MM subsets and muscularis polymorphonuclear cells (PMNs) from tamoxifen*-* and 1×DSS- or water-treated *Ccr2*^ER-Cre^*R26*-STOP^fl/fl^tdTomato mice at week 5, quantified by FC. (k) Bulk RNA-seq heatmap showing expression of MM Core Signature genes by monocyte-derived MM subsets from the same dataset as in (g). All graphs show mean ± SEM from 1 representative experiment. Statistical analyses: Unpaired Student’s t-test and two-way ANOVA for multiple comparisons, *P≤0.05, **P≤0.01, ***P≤0.001, and ****P≤0.0001.

Tissue macrophages differ in their developmental origins and evolve either from embryonic progenitors like brain microglia or from bone-marrow progenitors^24^. Intestinal mucosal macrophages develop mostly from monocytes both at steady state and inflammation^23,25–27^, whereas MMs are of a mixed origin in early postnatal life, and up to 20% remain embryonically derived in the normal adult intestine^22,23^. The origins of MMs in colitis have not been studied. To establish the source of MMs we first sought to establish their heterogeneity. Muscularis externa from the normal adult mouse colon was transcriptionally profiled at a single-cell level using 10X Genomics. We identified 20 cell clusters of 13 distinct cell types (in total 34,417 cells) including clusters of 6,929 MMs and 1,548 myenteric neurons, thus creating a comprehensive dataset to study MMs and their interaction with enteric neurons (Extended Data Fig. 2a, Table 1). Cell cluster analysis revealed significant cellular heterogeneity of MMs that were subdivided into five major clusters of *Csf1r*^+^*Flt3*^−^ cells (Extended Data Fig. 2b,c). Slingshot analysis of five MM clusters predicted the M1 cluster that co-expressed the highest levels of monocyte-specific markers *Ccr2* and *Ly6c2*, as well as fibronectin (*Fn1*), to be an immediate monocyte descendant and the last common ancestor for other MM clusters (Fig. 3e, d, Extended Data Fig. 2c). However, the M5 cluster expressed the markers of self-maintaining monocyte-independent macrophages *Lyve1*, *Timd4* and *Folr2*^28^. Using an earlier gene array dataset of intestinal macrophages^8,29^ we established the MM core gene signature, i.e., the list of genes with functions relevant to ENS homeostasis that were expressed more in MMs than in mucosal macrophages (Extended Data Fig. 2d). The MM core signature genes were differentially expressed by specific MM clusters. Some genes were highly expressed by M1 cluster, while other genes, particularly growth factors *Bmp2*, *Kitl*, *Igf* and BMP2 kinase *Bmp2k*, were most highly expressed by M4 and M5 clusters, suggesting the functional division of labor among MM clusters. Because M4 was the largest MM cluster and M1 and M4 clusters were predicted to be developmentally linked and monocyte-related, we applied CellChat analysis to identify and compare ligand-receptor pair interactions between M1 or M4 clusters and myenteric neurons. Although the ligand-receptor pairs for M1 and M4 clusters overlapped, some ligand-receptor pairs were unique to either M1 (e.g., *Sema3c*-*Plxnd1* and *Fn1*-*Cdc4*) or M4 (e.g., *App*-*Cd74* and *Gas6*-*Mertk*) clusters, suggesting that following recruitment into the myenteric plexus, monocytes engage in interactions with enteric neurons and undergo differentiation into mature tissue macrophages, acquiring distinct neuron-associated functions during this transition.

Next, we established the changes in MM heterogeneity during colitis. Our prior studies showed that MMs are a dominant immune cell type in the normal myenteric plexus^8^. The MM dominance remained after colitis induction, although the numbers of muscularis polymorphonuclear cells (M-PMNs) and muscularis T cells (M-T cells) also increased but to a lesser extent (Extended Data Fig. 3a-c). We identified five major MM populations based on their expression of monocyte/macrophage-specific markers (*Ccr2*-RFP, Ly6c, *Cx3cr1*-GFP, MHCII) that we referred to as CCR2^+^ MM1 (CCR2^+^, MHCII^−^, mostly Ly6c^+^ and CX3CR1^low^(lo)), CCR2^+^MHCII^+^ MMs further divided into Ly6C^+^ MM2 and Ly6c^−^ MM3, MHCII^+^ MM4 (CCR2^−^, Ly6c^−^, MHCII^+^, mostly CX3CR1^high^ (hi)) and CCR2^−^MHCII^−^ MM5 (Ly6C^−^, mostly CX3CR1^lo^) (Fig. 3f, Extended Data Fig. 3d). To link the scRNAseq M1-5 clusters to MM1-5 subsets we isolated and transcriptionally profiled the most numerous MM2, MM3, MM4, MM5 subsets from normal colon, and MM1, MM2, MM3 and MM4 subsets from inflamed colon using four-way FACS and bulk RNAseq (Fig. 3f). Normal (water) MM2, MM3, MM4 and MM5 subsets most corresponded to M1, M2, M4 and M5 clusters, respectively, based on their expression of several MM cluster-specific genes. Quantitively, the CCR2^+^ MM1 subset was negligible in the control group but expanded 10-fold upon colitis; the CCR2^+^MHCII^+^ MM2/MM3 subsets doubled whereas the MHCII^+^ MM4 and CCR2^−^MHCII^−^ MM5 subsets did not increase upon colitis (Fig. 3h). Although the monocyte marker Ly6c did not cover 100% of newly recruited CCR2^+^ MMs, changes in Ly6c^+^ and Ly6c^+^MHCII^+^ MM counts upon colitis positively correlated with increases in CCR2^+^ and CCR2^+^MHCII^+^ MMs (Extended Data Fig. 3e,f). Therefore, for further experiments where the use of *Ccr2*-RFP transgene was not feasible, *Ccr2*-RFP was replaced with the Ly6c antibody.

Confocal microscopy of the colonic muscularis externa confirmed that CCR2^+^ MMs expanded within the myenteric plexus during colitis (Figure 3i). In contrast, CD3^+^ T cells shown to participate in the killing of enteric neurons infected with neurotropic viruses^30^ were rare in the myenteric plexus of both control mice and mice with colitis (Extended Data Fig. 3g). Collectively, our analysis showed that colitis leads to a substantial expansion of monocyte-derived CCR2^+^ MMs.

To quantify the contribution of monocytes to the total MM pool, we fate-mapped monocyte-derived MMs at steady state and upon colitis in *Ccr2*^ER-Cre^*Rosa26*-STOP^fl/fl^tdTomato mice continuously treated with tamoxifen. Four days after beginning the tamoxifen treatment, nearly 100% of blood Ly6c^+^ inflammatory monocytes were tdTomato^+^ regardless of colitis (Extended Data Fig. 3h). Four and half weeks (33 days) later over 90% of Ly6c^+^ MMs and ∼75% of Ly6c^−^ MMs were labeled with tdTomato, i.e., they were mostly monocyte-derived. The rate of MM labeling with tdTomato was only slightly lower in control mice. In contrast, the rate of tdTomato^+^ cells among Ly6c^−^MHCII^−^ MMs was only 10% in control mice and 40% in DSS mice, suggesting their low turnover rate and/or low input from monocytes (Fig. 3l). Consistently, the scRNAseq M5 cluster and CCR2^−^MHCII^−^ MM5 subset that correspond to Ly6c^−^MHCII^−^ MMs expressed genes associated with monocyte-independent macrophages (*Lyve1*, *Timd4*, *Folr2*).

To understand the functional outcome of monocyte recruitment and MM expansion during colitis, we identified differentially expressed genes (DEGs) between normal (water MM2, MM3, MM4) and inflamed (DSS MM1, MM2, MM3, MM4) monocyte-derived MM subsets. While some ENS-homeostasis-related genes were reduced or unchanged in response to colitis, other DEGs with functions relevant to ENS homeostasis, neurodegeneration, inflammation and immune regulation were upregulated in a subset-specific manner that could not fit the definitions of “M1” pro-inflammatory and “M2” anti-inflammatory macrophage responses (Fig. 3k, Extended Data Fig. 3 i,j, Tables 2, 3, 4).

Taken together, we found that mucosal inflammation caused by DSS treatment leads to extensive monocyte recruitment into the ENS microanatomical compartment outside of the mucosa where monocytes sequentially differentiate into functionally distinct CCR2^+^ and CCR2^−^ENS-associated MMs and acquire microanatomical compartment-specific tissue-remodeling and microglia-like properties, likely to clear dying, repair injured and support newly born neurons.

### Monocyte-derived MMs recruited by enteric neurons via the CCL2/CCR2 axis facilitate ENS remodeling

To validate some of our transcriptional predictions, we assessed the dynamic changes in neuron-MM interactions during the active and receding stages of colitis. Mice were treated with one cycle of DSS (to have a better-controlled system) or water, and the interactions between MMs (MHCII^+^) and myenteric neurons (Hu^+^) were quantified by two-photon microscopy. Like in prior experiments, the initial loss of neurons observed at week 1 was followed by the recovery of their number (Fig. 4a,b). A significant increase in MM number was also observed but the peak of MM expansion (week 2) was delayed as compared to the peak of neuronal loss (week 1), suggesting that MM expansion is secondary to the initial neuronal loss. At the peak of MM expansion, there were increased contacts between MMs and neurons and increased neuronal (Hu) signal inside MMs (MHCII) reflecting MM recruitment into the myenteric ganglia and uptake of neurons by MMs (Fig. 4a,b). Thus, these results propose that MMs are recruited in response to inflammation-activated/damaged neurons.

**Figure 4.**
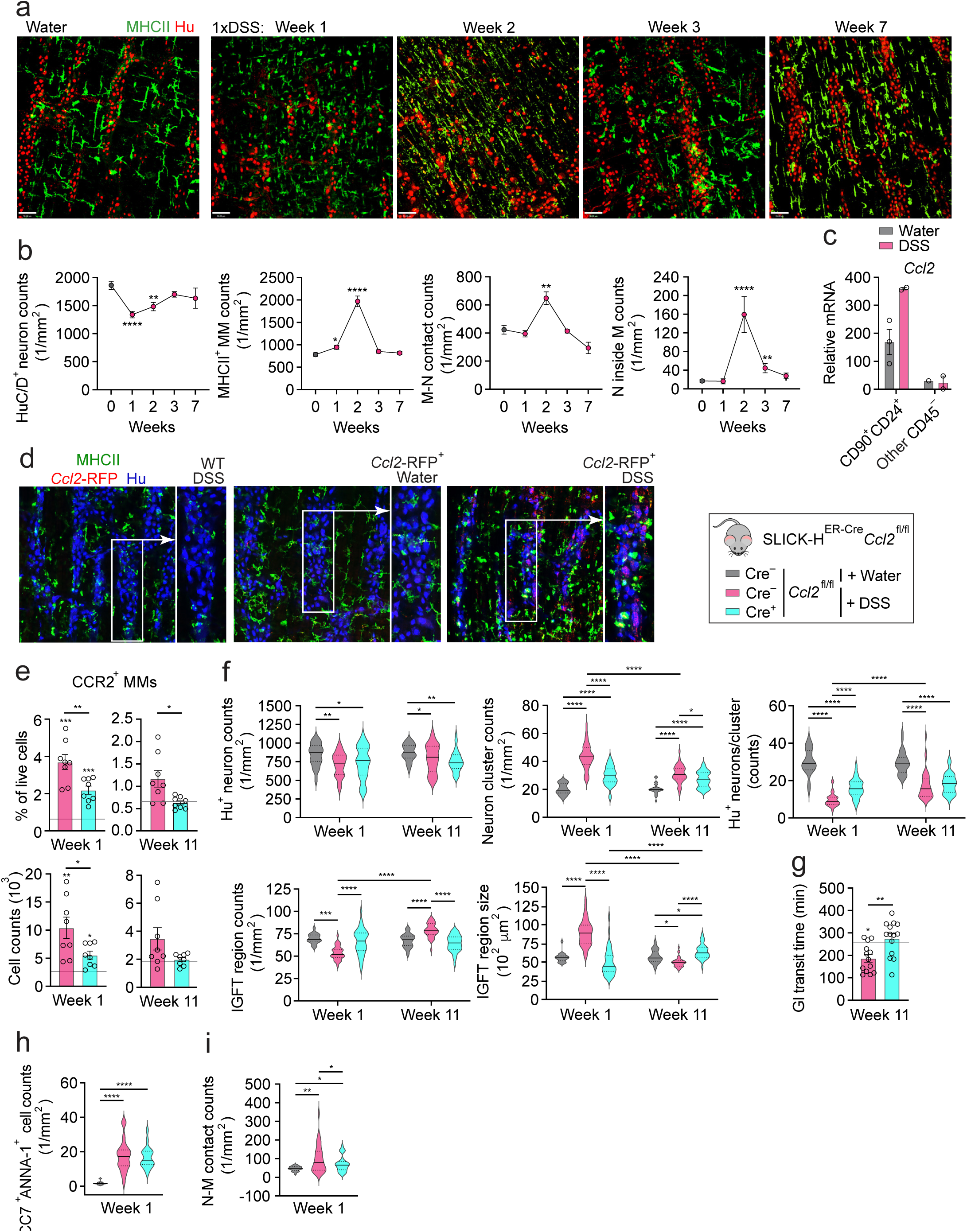
Enteric neuron-derived CCL2 facilitates pathological myenteric plexus remodeling and PI-GID by excessive recruitment of CCR2^+^ monocytes in response to colitis. (a) Multiphoton images of colonic myenteric plexus from 1×DSS- or water-treated mice. Scale bars, 50 μm. (b) MMs, myenteric neurons and their interactions quantified by multiphoton microscopy in the experiment described in (a). (c) *Ccl2* expression by qPCR in CD90^+^CD24^+^ myenteric neurons and other CD45^−^ stromal cells FACS-purified from colonic muscularis of 1×DSS- or water-treated mice at week 1. (d) Confocal images of colonic myenteric plexus from 1×DSS- or water-treated *Ccl2*-RFP^fl/fl^ and WT mice at week 1, stained with MHCII (MMs), tdTomato (*Ccl2*-RFP) and Hu (neuronal somas) antibodies. Scale bars, 100 μm. (e) Percentage of total viable cells and absolute cell counts of colonic CCR2^+^ MMs in tamoxifen- and 3×DSS-treated SLICK-H^ER-Cre^*Ccl2*^fl/fl^ and *Ccl2*^fl/fl^ littermates, quantified by FC. Baseline shows average value for tamoxifen- and water-treated *Ccl2*^fl/fl^ littermates (n=5-6). *P* values above each column show statistical differences over baseline. (f) Structural changes in the myenteric plexus from mice described in (e) quantified by confocal microscopy. Data are shown xas a distribution of pooled values per group. (g) GI transit in mice described in (e) (baseline n=9). (h) Counts of cleaved caspase 7 (CC7)^+^ANNA-1^+^ colonic myenteric neurons in mice described in (e), quantified by confocal microscopy. (i) Interactions (contacts) between MHCII^+^ MMs and myenteric Hu^+^ neurons in mice described in (e), quantified by confocal microscopy. All graphs show mean ± SEM from 1 representative experiment. Statistical analyses: Unpaired Student’s t-test and two-way ANOVA for multiple comparisons, *P≤0.05, **P≤0.01, ***P≤0.001, and ****P≤0.0001.

Next, we sought to identify an ENS-associated cell type responsible for recruiting CCR2^+^ monocytes into the ENS niche. In the myenteric plexus, only enteric neurons can directly access the mucosa through projected axons^31,32^. We therefore hypothesized that biological changes in enteric neurons should be most upstream to other changes in the microanatomical compartment of the myenteric plexus, and enteric neurons are the most likely cell type to recruit MMs. In agreement with our hypothesis, FACS-purified myenteric neurons but not other stromal cells expressed CCR2 ligand *Ccl2* at baseline and upregulated it further upon colitis (Fig. 4c). We confirmed CCL2 expression by myenteric neurons in the intact tissue by taking advantage of RFP expression under the control of *Ccl2* promoter (*Ccl2*-RFP) in *Ccl2*^fl/fl^ reporter mice (Fig. 4d). In colitis, the extent of neuronal CCL2 response in different ganglia varied, possibly reflecting heterogeneity of mucosal inflammation in different colonic regions that these neurons innervate. Other cell types that occasionally expressed CCL2 were MMs and non-neuronal inter-ganglionic cells likely representing enteric glia (Fig. 4d).

To prove the existence of the neuron-to-monocyte CCL2-CCR2 signaling axis, we conditionally depleted *Ccl2* in enteric neurons *in vivo*. We used SLICK-H^ER-Cre^ mice as a tamoxifen-inducible transgenic Cre line specific to neurons^33^. Currently, no other inducible Cre line exists that can target the majority of enteric neurons in an inducible manner. The inducible system was selected to avoid possible developmental defects caused by early MM loss^22^. The specificity of the SLICK-H^ER-Cre^ mouse model to myenteric neurons but not enteric glia was confirmed in SLICK-H^ER-Cre^*R26*-STOP^fl/fl^tdTomato mice (Extended Data Fig. 4a-c). Successful tamoxifen-induced *Ccl2* depletion in enteric neurons of SLICK-H^-ER-Cre^*Ccl2*^fl/fl^ (*Ccl2*^ΔNeu^) mice was also confirmed (Extended Data Fig. 5a). Next, *Ccl2*^ΔNeu^ mice and their *Ccl2*^fl/fl^ littermates were given 3×DSS and analyzed at early acute and late post-inflammatory time points (Fig. 4e). The depletion of *Ccl2* in enteric neurons reduced the number of recently recruited Ly6c^+^ MMs (for consistency referred to as CCR2^+^ MMs) both at early and late time points (Fig. 4f, Extended data Fig. 5b), while having no significant impact on muscularis PMN and T cells (Extended Data Fig. 5c,d). There was also no significant difference in mucosal inflammation at early inflammatory and late post-inflammatory time points as assessed by changes in the body weight, fecal lipocalin-2 levels and mucosal neutrophil counts (Extended Data Fig. 5e-g), and no difference in blood monocyte counts (Extended Data Fig. 4h) between the *Ccl2*-sufficient and deficient groups. These data demonstrate that during colitis enteric neurons recruit CCR2^+^ monocytes into the myenteric plexus by upregulating CCL2 production.

**Figure 5.**
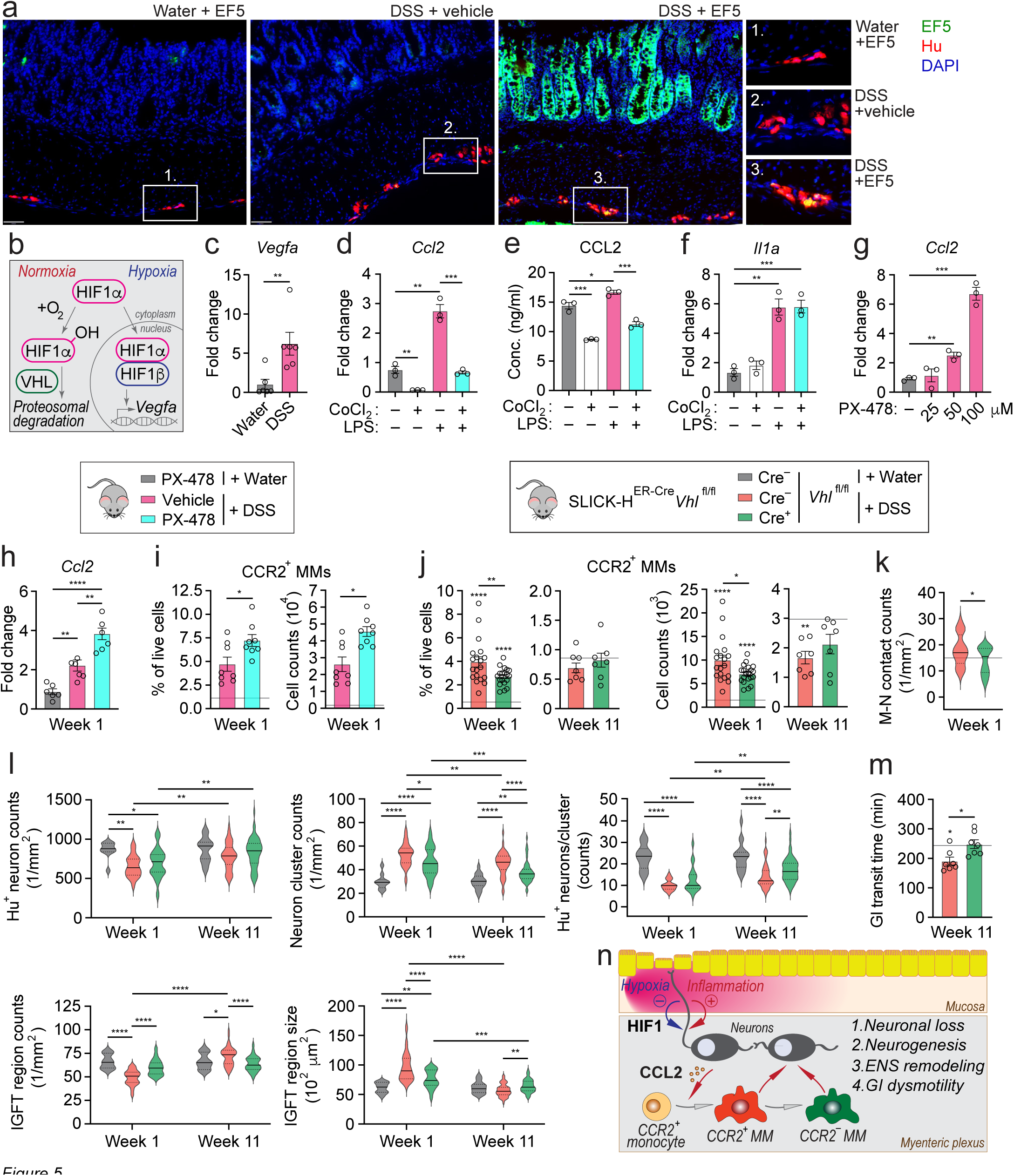
Enteric neuron-intrinsic HIF1α signaling limits CCL2-driven monocyte recruitment and protects from myenteric plexus remodeling and PI-GID. (a) Confocal images showing cross-sections of colons isolated from 1×DSS- or water-treated and EF5- or vehicle-injected WT mice at week 1, stained with EF5 (hypoxic cells) and Hu (neuronal somas) antibodies. Scale bars, 50 μm. (b) Graphical summary of hypoxia-inducible factor (HIF) pathway. (c) *Vegfa* expression by qPCR in colonic myenteric neurons FACS-purified from 1×DSS- or water-treated mice at week 1. (d) *Ccl2* expression by qPCR in cultured colonic myenteric neurons, treated with CoCl_2_, LPS or their combinations. Each data point is a well. (e) CCL2 concentration by ELISA in supernatants from neuronal culture described in (d). (f) *Il1a* expression by qPCR in neuronal culture described in (d). (g) *Ccl2* expression by qPCR in colonic neuronal culture treated with HIF1α inhibitor PX-478 or vehicle. (h) *Ccl2* expression by qPCR in CD90^+^CD24^+^ colonic myenteric neurons FACS-purified from 1×DSS- or water-treated WT mice daily injected with PX-478 or vehicle, analyzed at week 1. (i) Percentage of total viable cells and absolute cell counts of colonic CCR2^+^ MMs in mice described in (h), quantified by FC. Baseline shows average value for PX-478-injected mice on water (n=2). (j) Percentage of total viable cells and absolute cell counts of colonic CCR2^+^ MMs from tamoxifen- and 3×DSS-treated SLICK-H^ER-Cre^*Vhl*^fl/fl^ and *Vhl*^fl/fl^ littermates, quantified by FC. Baseline shows average value for tamoxifen- and water-treated *Vhl*^fl/fl^ littermates (n=6 and 5). (k) Interactions (contacts) between MHCII^+^ MMs and Hu^+^ myenteric neurons in mice described in (j) (baseline n=5). (l) Structural changes in myenteric plexus from mice described in (j) quantified by confocal microscopy. Data are shown as a distribution of pooled values per group. (m) GI transit in mice described in (j) (baseline n=5). (n) Graphical summary of HIF1α-CCL2 neuron-intrinsic regulatory pathway upstream of MM-driven ENS remodeling during colitis. All graphs show mean ± SEM from 1 representative experiment. Statistical analyses: Unpaired and paired Student’s t-test and two-way ANOVA for multiple comparisons, *P≤0.05, **P≤0.01, ***P≤0.001, and ****P≤0.0001.

*Ccl2*^ΔNeu^ mice provided a CCR2^+^ MM-deficient model for the follow-up functional studies. We assessed the role of CCR2^+^ MMs in colitis-associated ENS remodeling in the same cohort of mice with colitis as described above. We found that *Ccl2*^ΔNeu^ mice had diminished fragmentation of myenteric ganglia and less disrupted nerve fiber network at the early inflammatory phase and significantly less severe post-inflammatory remodeling of the myenteric plexus (Fig. 4f, Extended Data Fig. 6a). The difference in total myenteric neuron counts between the two colitis groups was not statistically significant although this was the least sensitive readout (see Methods). Consistent with reduced ENS remodeling, *Ccl2*^ΔNeu^ mice were protected from PI-GID (Fig. 4g).

Finally, we asked about the biological role of CCR2^+^ MMs in ENS remodeling. DSS and infectious colitis have been shown to increase programmed cell death of enteric neurons^12,17^. Because of the inflammatory gene signature of CCR2^+^ MMs during colitis (Extended data Fig 3j), we asked about their role in early neuronal apoptosis. We saw a significant increase in cleaved caspase-7 (CC7)^+^ apoptotic neurons in mice with colitis as compared to no colitis control, but there was no significant difference in the frequency of CC7^+^ neurons between *Ccl2*^ΔNeu^ and *Ccl2*^fl/fl^ groups with colitis (Fig. 4h, Extended Data Fig. 6b). In contrast, the physical interactions between MMs and myenteric neurons were reduced in the *Ccl2*^ΔNeu^ group (Fig. 4i). This evidence supports our earlier findings and lends further credence to our concept that MMs contribute to the cleanup of damaged neurons, creating conditions for neuronal repair rather than neuronal killing.

Collectively, we demonstrate that CCR2^+^ MMs are instrumental in the pathogenesis of PI-GID due to disproportionate ENS remodeling by MMs in response to colitis-associated ENS damage.

### Enteric neuron-intrinsic hypoxia signaling via HIF1α limits colitis-induced recruitment of monocyte-derived MMs and reduces PI-GID

Increased influx of inflammatory immune cells, their local proliferation and elevated rates of glycolysis during colitis increase oxygen consumption and result in suboptimal oxygen levels in the tissue termed “inflammatory hypoxia” as opposed to “physiological hypoxia” present in the normal colon^34^. We therefore hypothesized that enteric neurons and/or their projected axons located within the inflamed bowel regions become hypoxic. To test our hypothesis, mice with early DSS colitis and no colitis control mice were injected with a vehicle or the nitroimidazole compound EF5 which specifically labels hypoxic cells when injected *in vivo*. In DSS-treated mice, inflamed colonic regions with EF5^+^ (i.e., hypoxic) epithelium contained myenteric ganglia with EF5^+^ neurons (Fig. 5a). Thus, enteric neurons become hypoxic upon colitis.

The hypoxia-inducible factor (HIF) pathway regulates the expression of a broad range of genes that facilitate cell adaptation to hypoxia. HIFs function as heterodimers composed of an oxygen-regulated alpha subunit and a stably expressed beta subunit. HIF1α is ubiquitously expressed and in the brain, it was shown to regulate the neuronal response to hypoxia^35^. Under normoxia, cytoplasmic HIF1α is continuously subjected to hydroxylation and proteasomal degradation with the help of the von Hippel Lindau (VHL) E3 ubiquitin ligase complex. During hypoxia, HIF1α is not degraded and thus translocates into the nucleus where it dimerizes with HIF1β and binds to HIF-responsive elements in specific target genes^36^. Specifically, vascular endothelial growth factor α (VEGFα) is one of the key molecules positively regulated by HIF pathway^35,37^ (Fig. 5b). We, therefore, tested *Vegfa* expression in enteric neurons isolated from the myenteric plexus of mice at an early phase of DSS colitis and control mice, and found that colitis significantly upregulated neuronal *Vegfa* (Fig. 5c). Thus, mucosal inflammation triggers the hypoxia adaptation response in enteric neurons in addition to the inflammatory CCL2 response.

Interestingly, in primary fetal human astrocytes, HIF1α binding site was found in the CCL2 promoter^38^. We, therefore, hypothesized that HIF1 signaling regulates CCL2 expression in neurons during inflammation. To study this possible connection, we established cell cultures of myenteric neurons isolated from adult mice^39^ and treated them with cobalt chloride (CoCl_2_), a known stabilizer of HIF1α and activator of HIF signaling^40^, and thus modeled chemical hypoxia *in vitro*. To confirm that our *in vitro* hypoxia system is HIF1α dependent, cultured myenteric neurons were treated with CoCl_2_ in the presence of HIF1α inhibitor PX-478^41^, and *Vegfa* expression was quantified by qPCR. CoCl_2_ treatment upregulated *Vegfa* expression, but when combined with PX-478, *Vegfa* expression was reduced (Extended Data Fig. 7a).

Using this validated *in vitro* model system, we examined the role of the HIF pathway on *Ccl2* expression. Myenteric neurons cultured in the presence or absence of CoCl_2_, were treated with pro-inflammatory stimuli LPS or IL1β, both known as positive regulators of CCL2 expression in other cell types^42,43^. We found that like primary enteric neurons (Fig. 4c), cultured adult enteric neurons expressed detectable levels of *Ccl2* at baseline (Extended Data Fig. 7b) and significantly upregulated *Ccl2* expression upon LPS or IL1β treatment. Triggering the HIF pathway with CoCl_2_ reduced *Ccl2* expression both at baseline and upon stimulation (Fig. 5d, Extended Data 7c,d). Similar results were found for CCL2 protein expression (Fig. 5e). CoCl_2_ treatment did not impact cell viability (Extended Data Fig. 7e) or significantly change baseline and induced *Il1a* (Fig. 5f, Extended Data Fig. 7f) and *Csf1* (Extended Data Fig. 7g) expression, but it did induce *Vegfa* expression in the same cell cultures (Extended Data Fig. 7h). Furthermore, treatment with HIF1α inhibitor PX-478 upregulated *Ccl2* expression in cultured enteric neurons (Fig. 5g, Extended Data 7i). Collectively, our *in vitro* studies show that the HIF pathway negatively regulates CCL2 chemokine expression in enteric neurons.

To test the role of HIF pathway on CCL2-CCR2 enteric neuron-to-MM axis *in vivo*, we treated mice with PX-478 or a vehicle control along with DSS and analyzed them at an early time point. We found that enteric neurons from PX-478-treated mice with colitis upregulated more *Ccl2* (Fig. 5h) and had more CCR2^+^ MMs (Fig. 5i, Extended Data Fig. 8a) as compared to the vehicle-treated mice with colitis. In contrast, there was no difference in muscularis PMN numbers (Extended Data Fig. 8b) and mucosal inflammation (Extended Data Fig. 9a-c) between the groups. These results support our *in vitro* findings and show that the HIF pathway negatively regulates CCL2 expression in enteric neurons *in vivo* and attenuates CCR2^+^ monocyte recruitment into the myenteric plexus.

Because HIF1α inhibition with PX-478 is not restricted to enteric neurons, we turned to Cre/loxP transgenic mouse approaches for modulating HIF signaling in enteric neurons. To block HIF signaling in enteric neurons that would thus mimic PX-478 treatment, we generated transgenic SLICK-H^ER-Cre^*Hif1a*^fl/fl^ mice that deplete HIF1α specifically in neurons when treated with tamoxifen (*Hif1a*^ΔNeu^ mice). *Hif1a*^ΔNeu^ mice are expected to express more CCL2 in enteric neurons. Similar to PX-478-treated mice, *Hif1a*^ΔNeu^ mice with active colitis had more CCR2^+^ MMs as compared to Cre^−^ mice, i.e., they recruited more monocytes into the myenteric plexus (Extended Data Fig. 8c,d and 10a). To test the effect of LPS and IL1β signaling used as CCL2 mimetics in the *in vitro* system, we developed SLICK-H^ER-Cre^*Myd88*^fl/fl^ mice that lose MyD88 (an adaptor molecule downstream of both inflammatory pathways) in neurons when treated with tamoxifen (*Myd88*^ΔNeu^ mice). *Myd88*^ΔNeu^ mice are expected to have reduced CCL2 expression in enteric neurons. Consistently, *Myd88*^ΔNeu^ mice with active colitis had fewer CCR2^+^ MMs as compared to Cre^−^ mice, i.e., they recruited fewer monocytes into the myenteric plexus (Extended Data Fig. 8f,g). Finally, to translate the *in vitro* HIF1α stabilization by CoCl_2_ into an *in-vivo* system, we generated SLICK-H^ER-Cre^*Vhl*^fl/fl^ mice that deplete a negative regulator of HIF signaling *Vhl* in neurons upon tamoxifen treatment (*Vhl*^ΔNeu^ mice). *Vhl*^ΔNeu^ mice are expected to have suppressed CCL2 expression due to more stabilized HIF1α available to trigger HIF signaling. As predicted, *Vhl*^ΔNeu^ mice with active colitis had fewer CCR2^+^ MMs as compared to Cre^−^ mice, i.e., they recruited fewer monocytes into the myenteric plexus (Fig. 5j, Extended Data Fig. 8i). Neither *Hif1a*^ΔNeu^, *Myd88*^ΔNeu^ nor *Vhl*^ΔNeu^ mice had significant differences in muscularis PMN numbers (Extended Data Fig. 8e,h,j) and mucosal inflammation (Extended Data Fig. 9d-i), confirming that the effect that we see on MMs is myenteric plexus-restricted and neuron-driven. Together, these results demonstrate that enteric neurons employ HIF signaling to negatively regulate inflammation-mediated recruitment of monocytes to the ENS. Further, they underscore the role of HIF signaling as a neuron-intrinsic adaptation program that limits monocyte recruitment to the myenteric plexus likely to prevent further escalation of tissue hypoxia and subsequent cell damage.

Finally, we examined the effect of colitis-associated HIF signaling on ENS remodeling leading to PI-GID. Confocal microscopy showed an increase in contacts between MMs and myenteric neurons and increased fragmentation of ganglia in *Hif1a*^ΔNeu^ mice with colitis (Extended Data Fig. 10a-c), consistent with our finding that ENS remodeling is driven by MMs. Conversely, *Vhl*^ΔNeu^ mice with hyperactive neuronal HIF signaling and reduced monocyte recruitment had fewer contacts between MMs and myenteric neurons and were partially protected from colitis-associated ENS remodeling (Fig. 5k,l, Extended Data Fig. 10d). Finally, *Vhl*^ΔNeu^ mice with a history of colitis had normal GI transit in contrast to Cre^−^ mice (Fig. 5m). Thus, hyperactivation of hypoxia-induced HIF signaling by VHL removal protects from pathological ENS remodeling and subsequent PI-GID (Fig. 5n).

### Human ENS is remodeled in patients with IBD

To test if ENS is remodeled in patients with IBD, we analyzed intestinal innervation in the inflamed and non-inflamed colon regions surgically resected from the same patients with CD and in non-IBD non-inflamed controls. A significant increase in the density of HLA-DR^+^ cells was found in the mucosa of patients with CD as compared to non-IBD control, with the highest HLA-DR^+^ cell density in inflamed CD regions, consistent with the degree of inflammation determined macroscopically. Similarly, the density of βIII-Tubulin^+^ nerve fibers was higher in CD samples as compared to non-IBD control, and the density in the inflamed CD regions was higher than in non-inflamed CD regions. The neuron density in the enteric ganglia was also significantly increased in the inflamed CD samples (Fig. 6a-c). Collectively, these data in IBD patients are consistent with the evidence of substantial ENS remodeling in mice with colitis.

**Figure 6.**
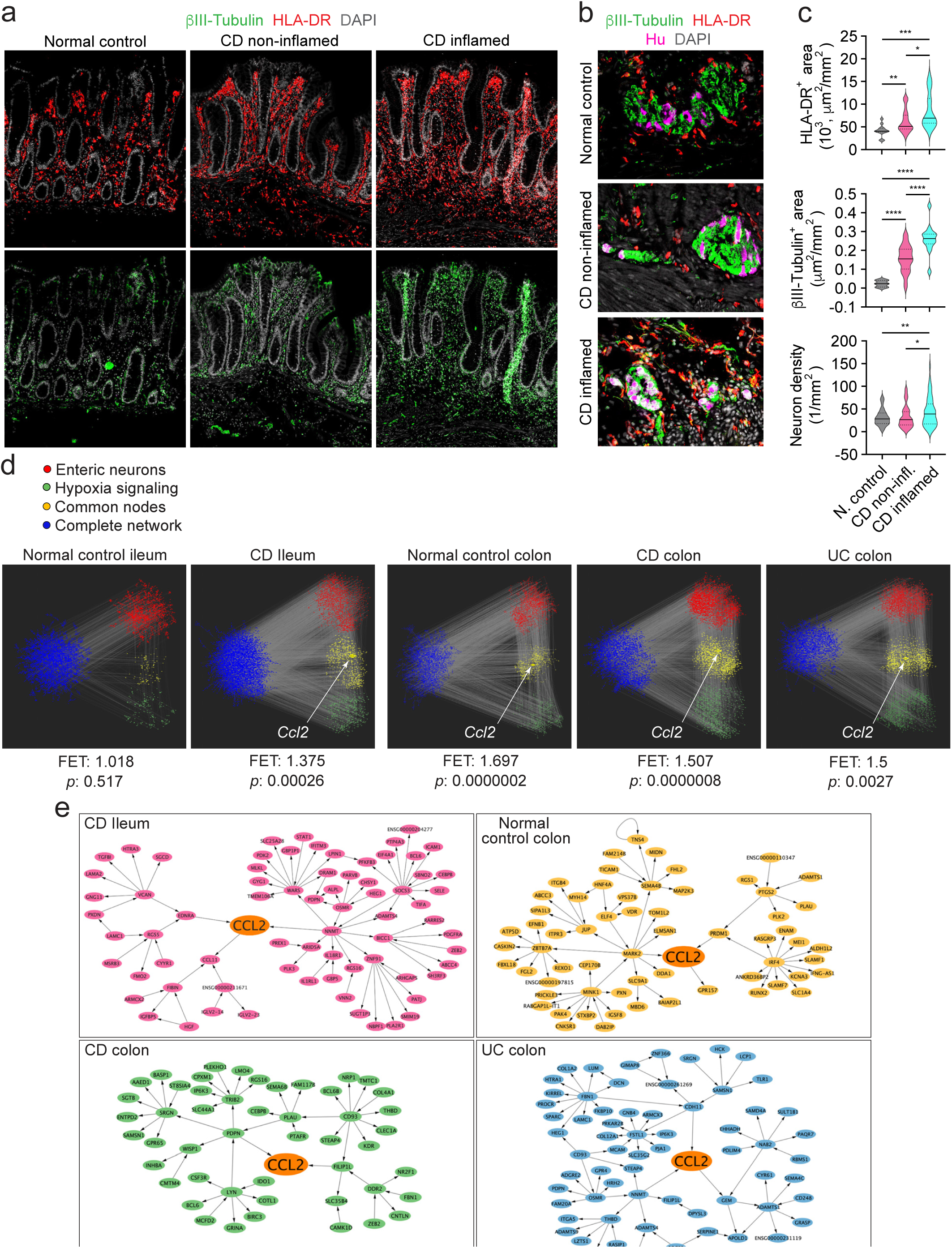
Human ENS is remodeled in patients with IBD through similar mechanisms. (a) Confocal images of large bowel mucosa in transmural specimens surgically resected from patients with colorectal cancer (CRC, n=3, normal adjacent region) and Crohn’s disease (CD, n=3, non-inflamed control and inflamed regions), stained with ýIII-Tubulin (nerve fibers), HLA-DR (including macrophages) and Hu (neuronal somas). Scale bars, 200 μm. (b) Confocal images of large bowel muscularis from the same tissue cross-sections as in (a). Scale bars, 100 μm. (c) HLA-DR^+^ cells (inflammation), ýIII-Tubulin^+^ nerve fibers (innervation) and Hu^+^ neuronal soma density in ýIII-Tubulin^+^ ganglia in the experiment described in (a) and (b), quantified by confocal microscopy. Data shown as a distribution of pooled values per group. (d) Hypoxia and inflamed enteric neuron sub-networks constructed in the following Bayesian networks (BN): control ileum, CD ileum, control colon, CD colon and ulcerative colitis (UC) colon. (e) The CCL2 sub-networks comprised from identifying the CCL2 node and extending out from CCL2 to include all directly connected nodes within three path lengths in the IBD and control Bayesian networks described in (d).

Subsequently, we investigated whether the gene associations identified in our mouse study are also observed in patients with IBD. We leveraged Bayesian networks used for discovering relations between genes, constructed from biopsies collected from the ileum and colon of IBD patients, including inflamed and non-inflamed tissue, and non-inflamed samples from routine screening as controls (Fig. 6d). These probabilistic networks infer the causal regulatory hierarchy of co-expressed genes in five specific Bayesian networks (BNs): control ileum, CD ileum, control colon, CD colon and UC colon^44,45^. To identify the homologous hypoxia sub-network in each network, we collated a hypoxia signature comprised of genes downstream of HIF signaling^46^ and CCL2/CCR2^42,43^ pathways derived from the literature. In projecting this hypoxia gene signature onto each network, we identified all gene network nodes overlapping between the signature and the network and included all nodes extending out two pathlengths from the shared signature and network nodes, to extract the largest connected subnetwork, comprising the hypoxia subnetwork of each network. Using a cell identity signature we generated for inflamed enteric neurons^47^, we tested fold enrichment in the hypoxia subnetwork relative to the full BN for each of the five networks. We found in the control ileum, there was no significant overlap between hypoxia and enteric neuron networks, however in the CD ileum, there was a mild significant overlap. As expected, the enrichment between enteric neurons and hypoxia response was greater in the normal colon than in the normal ileum^48^; however, the control colon (FET 1.697, *p*-value 0.0000002) had a slightly more significant enrichment than the CD colon (FET 1.507, *p*-value 0.0000008) or UC colon (FET 1.5, *p*-value 0.0027). CCL2 was not included in the control ileum hypoxia subnetwork, where the overlap was not significant, but CCL2 was present within the hypoxia subnetworks of the other four networks where the overlap with hypoxia and enteric neurons was significant.

When assessing the local network neighborhood structure of CCL2, which comprised all gene nodes directly connected within three path lengths of CCL2, the subnetworks encompassed pathways enriched in neurogenesis, remodeling, fibrosis, and hypoxia. There were several nodes shared across subnetworks, but most nodes were specific to each CCL2 subnetwork’s disease status and subset (Fig. 6e). We found that CCL2 subnetworks in disease BNs resolve processes related to hypoxia, fibroblast/neuronal interaction, angiogenesis, and smooth muscle remodeling. In contrast, in the control colon network, CCL2 is predicted to regulate anti-inflammatory genes, including TH17 regulation. The CCL2 IBD subnetworks include putative genes involved in the remodeling program such as: *ZEB1* (promotes the proliferation and differentiation of enteric neural precursor cells), FIBIN (secreted growth factor in neurons), *PDK2* (neuronal hypoxia response negatively correlated with macrophages), *MLKL* (involved in microglial neuronal nerve injury macrophage recruitment), *PDGFRA* (marker for neural fibroblasts that interact with macrophages) present in CD ileum; *DAB2IP*, *MARK2* (neurodegeneration), *GPR157* (regulates neuronal differentiation of glial progenitors), *ELMSANI* (regulator of neuronal differentiation), *TOMIL2* (neural degeneration), *SEMA4B* (neuron protection following injury via hypoxia signaling) present in control colon; *NRP1* (co-receptor for NGF that interacts with VEGF on endothelial cells and is inhibited by activation of AHR), *NR2F1* (involved in neuronal differentiation and fibroblasts), *SAMSN1* (neural stem cell proliferation and neuroprotection from hypoxic injury), *ENTPD2* (in colitis, ablation alters myenteric neuron numbers), *SGTB* (promotes neurite outgrowth and upregulation and is associated with neuronal apoptosis after neuroinflammation) present in CD colon; *DCN* (nerve injury), *SRGN* (amplifies microglial mediated neuroinflammation), *HTRA1* (neural crest migration and fibrosis), *KIRREL* (cell adhesion regulating neuronal interaction and angiogenesis), MCAM (may mediate a fibroblast-neuron axis and interacts with macrophages), *PRKAR2B* (neuroinflammation, hypoxia) present in UC colon. Collectively, these nodes may predict novel disease targets or biomarkers for further evaluation and prospective validation in follow-up studies for disease types presenting with enteric remodeling and PI-GID.

In summary, the findings in human IBD are in line with the cellular and molecular pathways found in our mouse studies.

## DISCUSSION

Our study established the pathogenesis of colitis-associated PI-GID. We revealed that prolonged colitis results in substantial structural ENS remodeling that is pathological in the context of PI-GID. ENS remodeling is driven by a combination of partial neuronal loss caused by increased incidence of programmed cell death and comparable gain of enteric neurons due to enhanced neurogenesis. Mucosal inflammation leads to the neurogenic CCL2-driven monocyte recruitment into the extra-mucosal myenteric plexus for the uptake and clearance of likely damaged neurons. However, newly formed monocyte-derived MMs promote ENS remodeling which we propose is a disproportionate ENS “repair” with PI-GID as its functional outcome. Finally, we identify the enteric neuron-intrinsic anti-inflammatory HIF-CCL2 axis that, when hyperactivated, restricts monocyte recruitment into the myenteric plexus and prevents PI-GID.

Prior studies have viewed MMs as sessile microglia-like macrophages of partial embryonic origin that switch their gene signature to a neuro-protective M2 phenotype in response to mucosal challenge^9,12,23^. Our study reveals that the extra-mucosal myenteric plexus represents a dynamic cellular compartment along with the intestinal mucosa, and monocytes known to infiltrate the inflamed mucosa early in the disease are also recruited in the myenteric plexus by activated enteric neurons. Once in the myenteric plexus, monocytes rapidly engage in functional interactions with enteric neurons and continue their differentiation trajectory into tissue MMs. The phenotype of newly recruited monocytes significantly differs from more differentiated monocyte-derived MMs, but both express pro-and anti-inflammatory genes along with genes important for ENS homeostasis and function suggesting division of labor with different contributions to ENS remodeling. Future studies will address the role of monocyte-derived MM-specific genes in ENS repair and remodeling.

In the myenteric plexus, only enteric neurons can directly access the mucosa through projected axons^31,32^ suggesting that they directly sense mucosal inflammation and transmit the signal to the myenteric plexus. This anatomical connection still needs to be proven experimentally, and the exact type of neurons involved in mucosal sensing of inflammation needs to be established.

HIF pathway has been shown to play a protective role in DSS colitis with an intrinsic role in the intestinal epithelium^49,50^. Combined with our findings of the protective role of HIF signaling in pathological ENS remodeling supported by extrapolative human data from patients with IBD, our study proposes that pharmacological bolstering of HIF pathway in patients with IBD could be used as a therapeutic strategy to prevent PI-GID and improve the overall gut health by restricting excessive ENS remodeling by inflammatory MMs.

### Statistical analyses

Statistical analysis was performed using GraphPad Prism 10 software. All data are presented either as mean ± SEM or as specified in the figure legends. Two-tailed paired or unpaired Student’s *t*-test was used to determine the statistical significance of differences when comparing means of 2 groups. Two-way analysis of variance (ANOVA) was used to determine the statistical significance when comparing more than 2 groups. Differences between groups were considered statistically significant when values of *p* ≤ 0.05.

---

**Figure S1.**
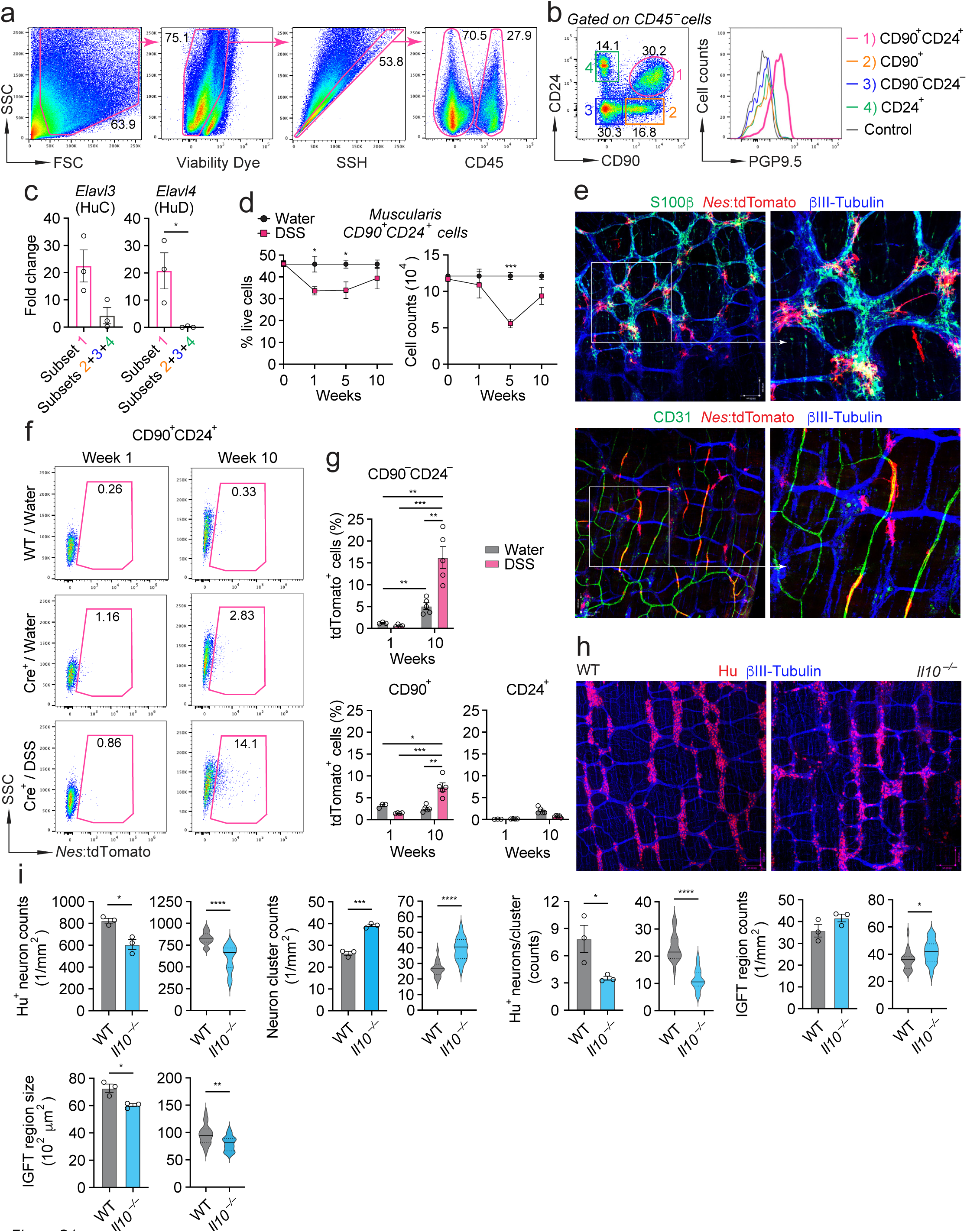
Colitis facilitates neurogenesis and structural remodeling of the myenteric plexus. (a) FC gating strategy to identify CD45^+^ immune cells (including MMs) and CD45^−^ stromal cells (including myenteric neurons) in colonic muscularis. (b) Intracellular expression of pan-neuronal marker PGP9.5 by four CD45^−^ subsets identified based on their co-expression of CD90 and CD24 shown by FC. (c) Relative expression by qPCR of pan-neuronal genes *Elavl3* (HuC) and *Elavl4* (HuD) by FACS-purified CD45^−^ subsets defined in (b). (d) Percentage of total viable cells (left) and absolute cell counts (right) of colonic muscularis CD90^+^CD24^+^ neurons in 3×DSS- or water-treated mice quantified by FC. (e) Confocal images of colonic myenteric plexus isolated from tamoxifen- and 1×DSS treated *Nes*^ER-Cre^*R26*-STOP^fl/fl^tdTomato (Cre^+^) mice on day 45 (6.5 weeks) and stained with antibodies to S100β (enteric glia) or CD31 (blood endothelial cells), tdTomato (newborn cells) and βIII-Tubulin (nerve fibers). Scale bars, 100 μm. (f) FC plots showing percentage of newly generated tdTomato^+^ cells among total CD45^−^ CD90^+^CD24^+^ myenteric neurons in colonic muscularis isolated from tamoxifen- and 3×DSS or water-treated *Nes*^ER-Cre^*R26*-STOP^fl/fl^tdTomato (Cre^+^) mice as compared to *R26*-STOP^fl/fl^tdTomato (Cre^−^) mice. (g) Percentage of newly generated tdTomato^+^ cells among CD45^−^ CD90^−^CD24^−^, CD90^+^ and CD24^+^ stromal cells in colonic myenteric plexus from mice described in (f), quantified by FC. (h) Confocal images of colonic myenteric plexus from 12-week-old *Il10*^−/−^ and WT mice stained with Hu and βIII-Tubulin antibodies. Scale bars, 100 μm. (i) Structural changes in myenteric plexus of mice described in (h) imaged by confocal microscopy. Data are shown as mean value per group (left) or as a distribution of pooled values per group (right). All graphs show mean ± SEM from 1 representative experiment. Statistical analyses: Unpaired Student’s t-test and two-way ANOVA for multiple comparisons, *P≤0.05, **P≤0.01, ***P≤0.001, and ****P≤0.0001.

**Figure S2.**
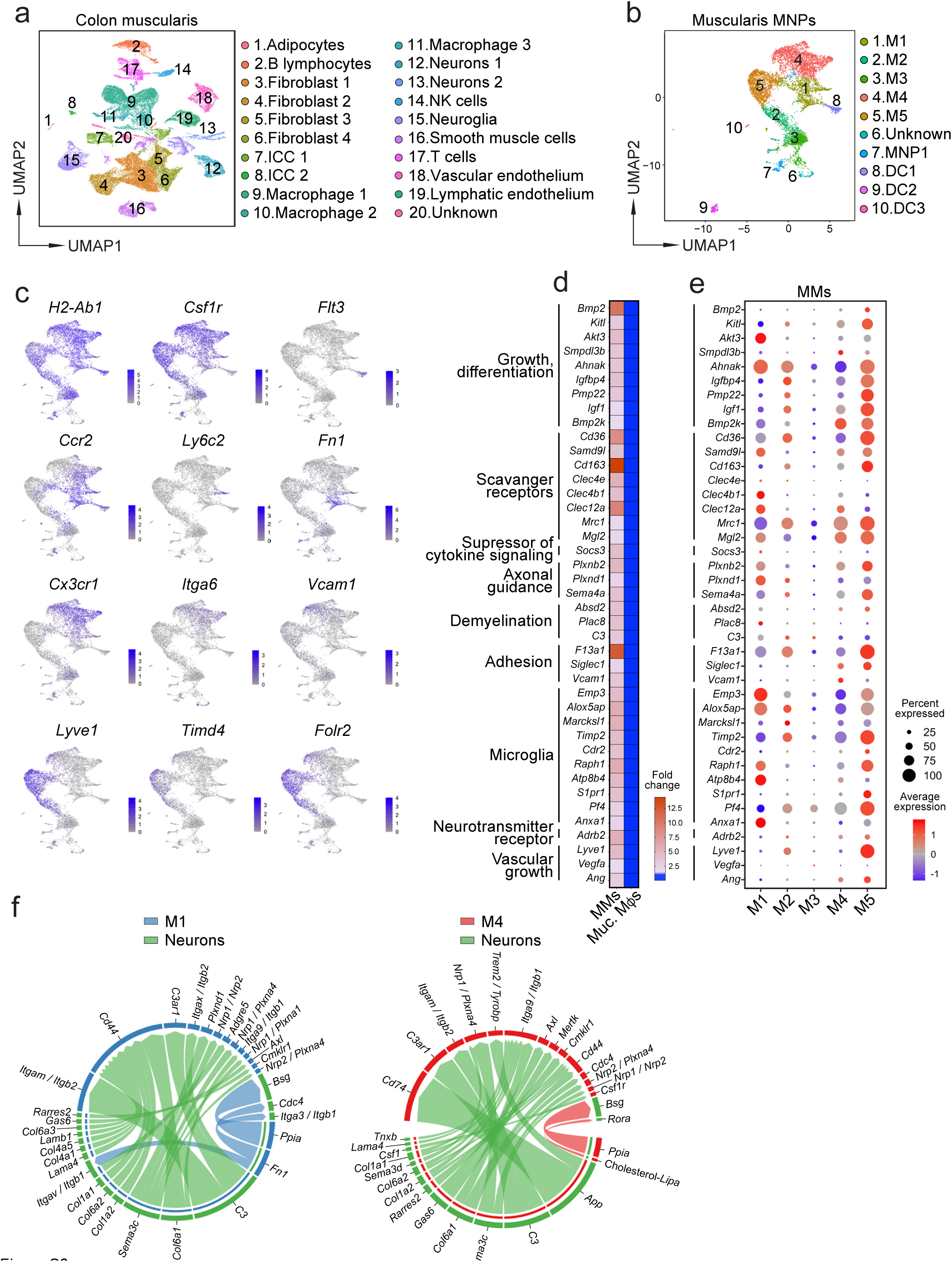
Transcriptomic analyses of colonic muscularis and MM subsets in the myenteric plexus at steady state. (a) UMAP clustering analysis of scRNA-seq dataset generated from colonic muscularis of normal adult WT mice. Cell type for each cluster annotated based on specific marker genes. (b) UMAP clustering analysis of all mononuclear phagocytes (MNPs) in the dataset described in (a). MM subset (M), dendritic cell (DC). (c) Normalized gene expression of 11 marker genes across muscularis MNPs identified in (b) (supplementary to Fig. 3e). (d) Heat map showing genes highly expressed in MMs as compared to mucosal macrophages (Muc. Mýs) in Immgen gene array dataset generated from normal small bowel MNPs. (e) ScRNA-seq dot heatmap showing expression of MM Core Signature genes from (d) across five MM clusters defined in (b). Color denotes the average expression level and size represents the percentage of cells expressing the indicated genes. (f) ScRNA-seq wheel plots showing top ligand-receptor pair interactions between colonic myenteric neurons and M1 (left) or M4 (right) monocyte-derived MM clusters predicted by CellChat analysis of the dataset described in (a).

**Figure S3.**
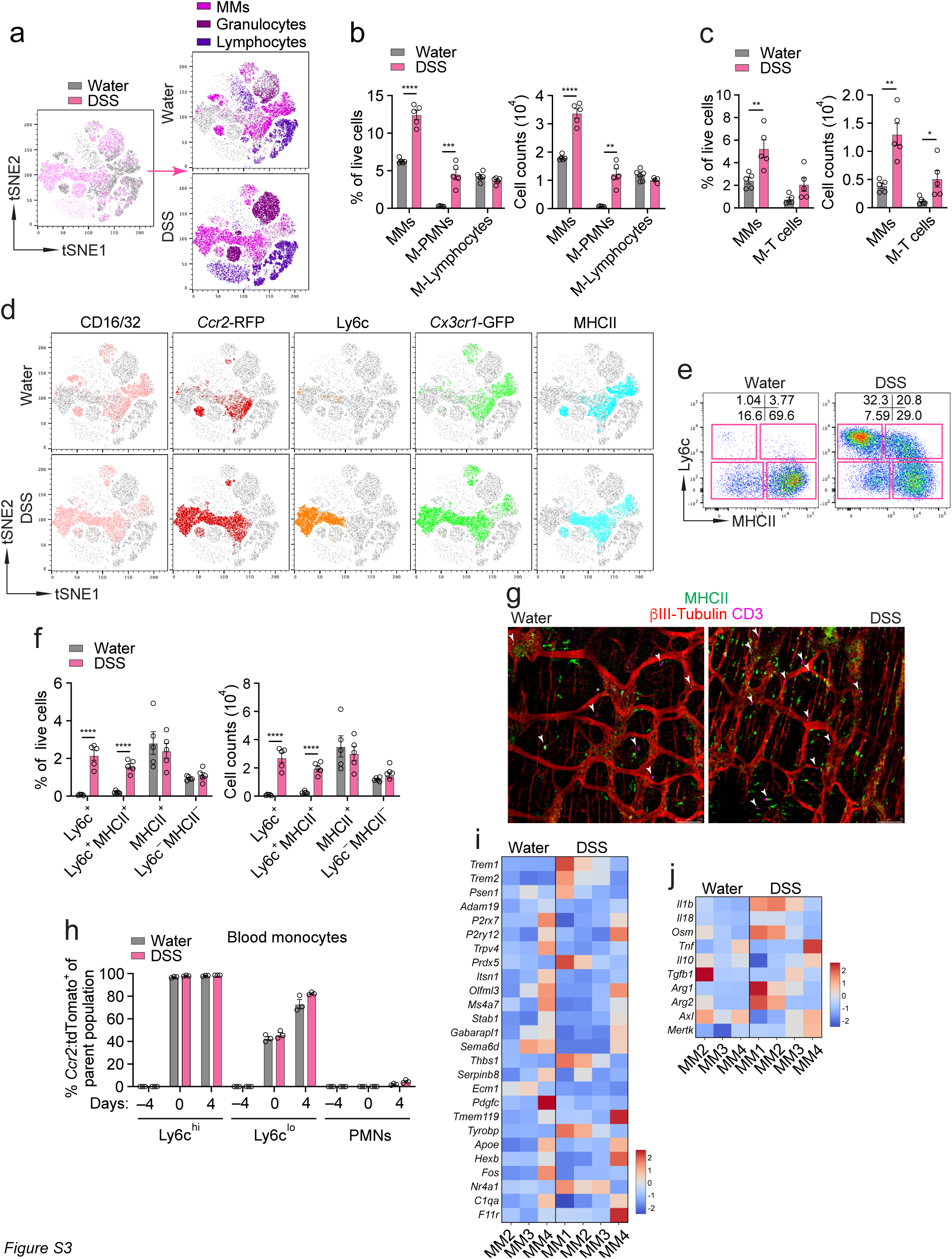
Monocyte-derived MMs are the dominant immune cell type expanding in the myenteric plexus in response to colitis. (a) FC tSNE plots showing immune cell landscape of colonic muscularis single-cell suspensions from 1×DSS- or water-treated *Ccr2*^RFP/+^*Cx3cr1*^GFP/+^ reporter mice at week 1. (b) Percentage of total viable cells (left) and absolute cell counts (right) of major immune cell types in the experiment described in (a), quantified by FC. (c) Percentage of total viable cells (left) and absolute cell counts (right) of MMs and muscularis T cells (M-T cells) in 1×DSS- or water-treated WT mice at week 1, quantified by FC. (d) FC tSNE plots showing the phenotype of colonic MM subsets in the experiment described in (a). (e) FC plots showing colonic MM subsets and their percentages based on Ly6c expression in 1×DSS- or water-treated *Ccr2*^RFP/+^*Cx3cr1*^GFP/+^ mice described in (a). (f) Percentage of total viable cells (left) and absolute cell counts (right) of MM subsets based on Ly6c expression in the experiment described in (a), quantified by FC. (g) Fluorescent microscopy images of colonic myenteric plexus from 1×DSS- or water-treated WT mice at week 1, stained with MHCII, βIII-Tubulin and CD3 antibodies. Scale bars, 75 μm. White arrows point at CD3^+^ T cells. (h) Percentage of tdTomato^+^ cells among Ly6c^hi^ and Ly6c^lo^ monocytes, and PMNs in the blood of tamoxifen- and 1×DSS or water-treated *Ccr2*^ER-Cre^*R26*-STOP^fl/fl^tdTomato mice, quantified by FC. (i) Bulk RNA-seq heatmap shows expression of brain microglia-specific genes across monocyte-derived MM subsets isolated from 1×DSS- or water-treated *Ccr2*^RFP/+^*Cx3cr1*^GFP/+^ mice at week 1. (j) Expression of inflammatory and anti-inflammatory genes across monocyte-derived MM subsets from the dataset described in (i). All graphs show mean ± SEM from 1 representative experiment. Statistical analyses: Unpaired Student’s t-test and two-way ANOVA for multiple comparisons, *P≤0.05, **P≤0.01, ***P≤0.001, and ****P≤0.0001.

**Figure S4.**
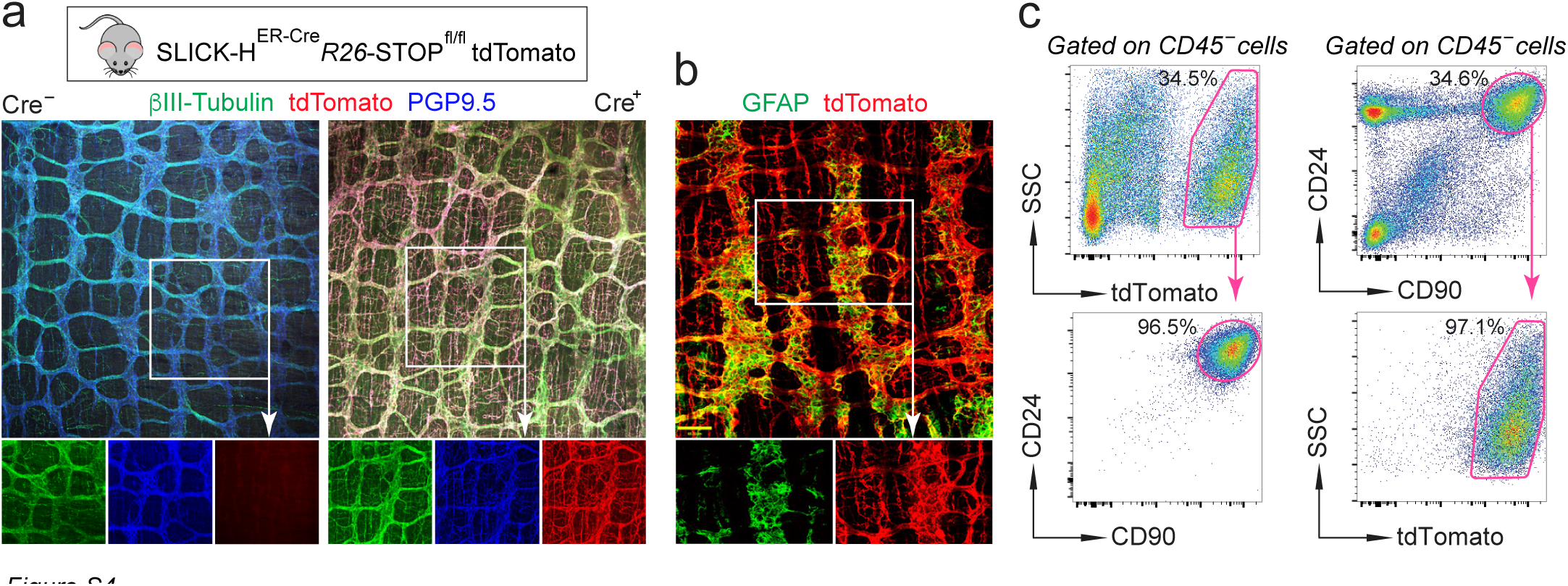
Tamoxifen-inducible SLICK-H^ER-Cre^ transgene specifically targets neurons in the colonic myenteric plexus. (a) Confocal images of colonic myenteric plexus from tamoxifen-treated SLICK-H^ER-Cre^*Rosa26*-STOP^fl/fl^tdTomato (Cre*^+^*) and control *Rosa26*-STOP^fl/fl^tdTomato (Cre^−^) mice, stained with βIII-Tubulin (nerve fibers), tdTomato (Cre-targeted cells) and PGP9.5 (whole neurons) antibodies. (b) Confocal images of colonic myenteric plexus from mice described in (a), stained with GFAP (enteric glia) and tdTomato (Cre-targeted cells) antibodies. Scale bars, 68 μm. (c) FC plots showing percentage of tdTomato^+^ cells among CD45^−^ CD90^+^CD24^+^ cells in colonic muscularis of tamoxifen-treated SLICK-H^ER-Cre^*R26*-STOP^fl/fl^tdTomato mice as compared to Cre^−^ controls.

**Figure S5.**
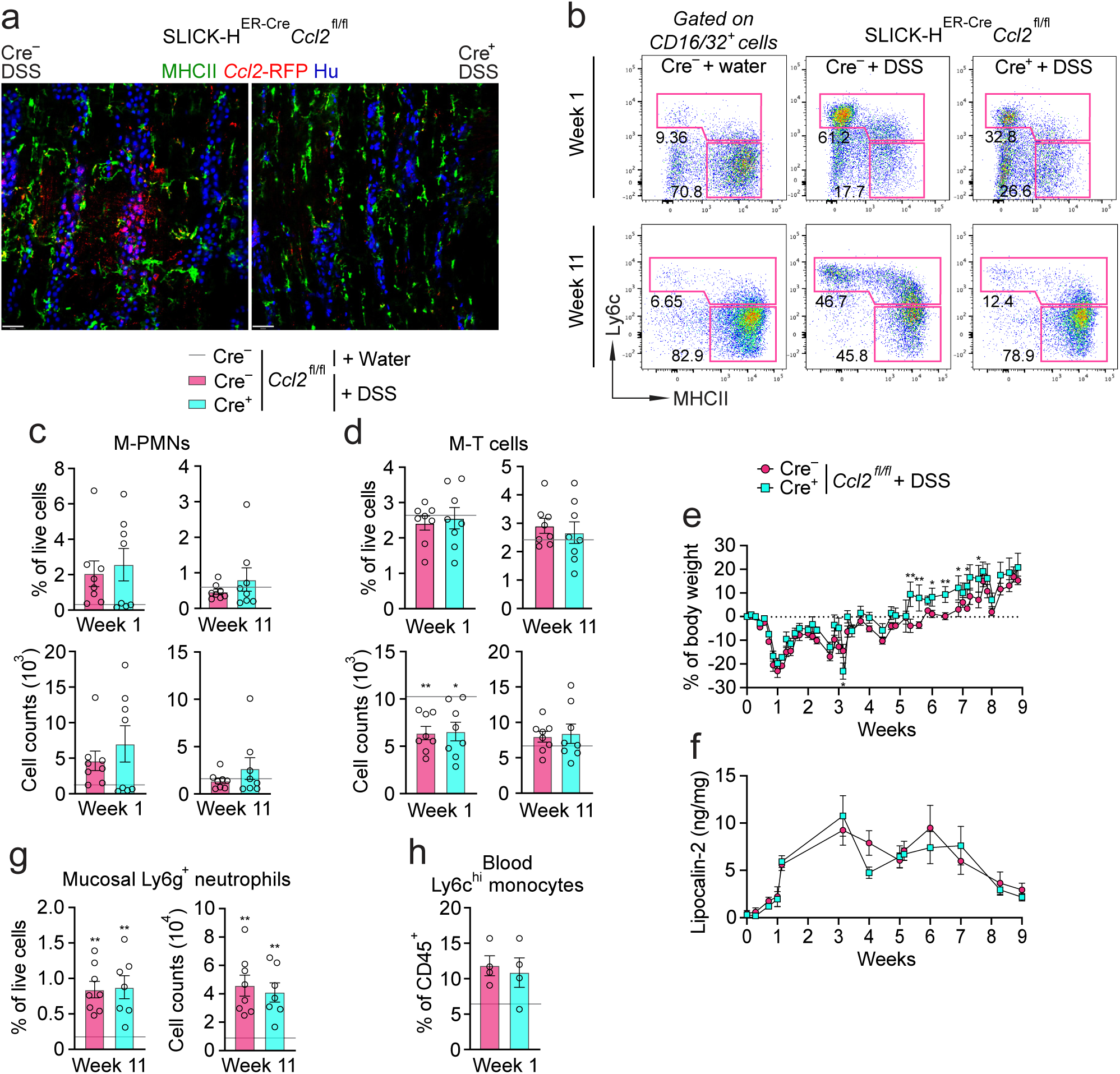
The efficacy of CCL2 depletion and extent of intestinal inflammation in SLICK-_HER-Cre*Ccl2*fl/fl mice._ (a) Confocal images of colonic myenteric plexus from tamoxifen- and 1×DSS-treated SLICK-H^ER-Cre^*Ccl2*-RFP^fl/fl^ (Cre^+^) and *Ccl2*-RFP^fl/fl^ (Cre^−^) littermates at week 1, stained with MHCII (MMs), tdTomato (*Ccl2*-RFP) and Hu (neuronal somas) antibodies. Scale bars, 50 μm. (b) FC plots showing proportions of CCR2^+^ (Ly6c^+^) and CCR2^−^ (Ly6c^−^) MM subsets and their percentages in colons of tamoxifen- and 3×DSS- or water-treated Cre^+^ and Cre^−^ littermates described in (a). (c) Percentage of total viable cells and absolute cell counts of colonic muscularis PMNs (M-PMNs) in mice described in (b), quantified by FC. Baseline shows average value for tamoxifen- and water-treated Cre^−^ littermates (n=6). (d) Percentage of total viable cells and absolute cell counts of colonic muscularis T cells (M-T cells) in mice described in (b), quantified by FC (baseline n=6). (e) Percentage of body weight change in tamoxifen- and 3×DSS-treated mice described in (b). (f) Fecal lipocalin-2 in tamoxifen- and 3×DSS-treated mice described in (b). (g) Percentage of total viable cells and absolute cell counts of colonic mucosal Ly6g^+^ neutrophils in mice described in (b), quantified by FC (baseline n=6). (h) Percentage of blood Ly6c^hi^ monocytes among total viable CD45^+^ blood cells in mice described in (b), quantified by FC (baseline n=3). All graphs show mean ± SEM from 1 representative experiment. Statistical analyses: Unpaired Student’s t-test and two-way ANOVA for multiple comparisons, *P≤0.05, **P≤0.01, ***P≤0.001, and ****P≤0.0001.

**Figure S6.**
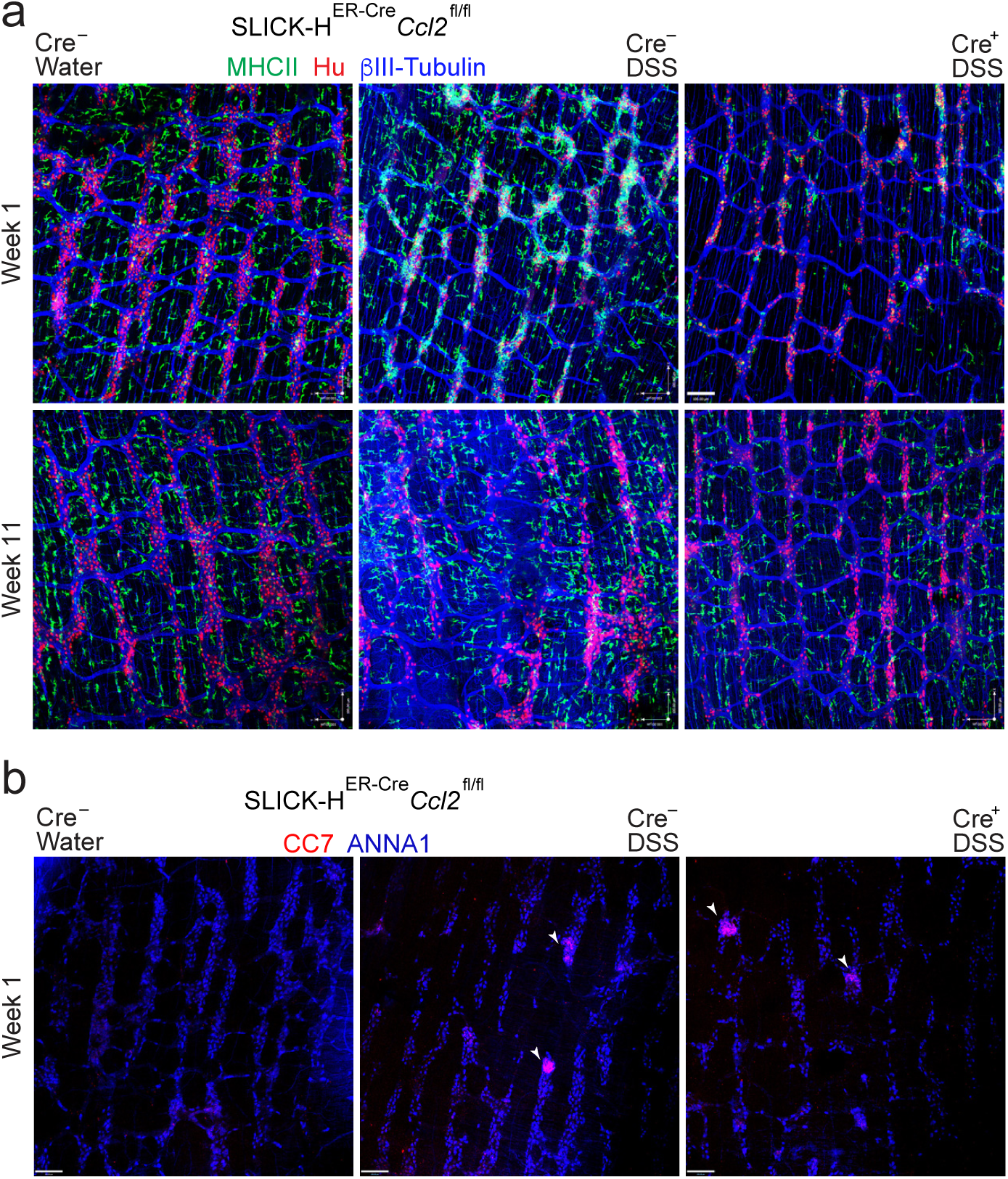
Monocyte-derived MMs recruited by enteric neurons via the CCL2/CCR2 axis facilitate myenteric plexus remodeling. (a) Confocal images of colonic myenteric plexus from tamoxifen- and 3×DSS- or water-treated SLICK-H^ER-*Cre*^*Ccl2*^fl/fl^ (Cre^+^) and *Ccl2*^fl/fl^ (Cre^−^) littermates, stained with MHCII (MMs), Hu (neuronal somas) and ýIII-Tubulin (nerve fibers) antibodies. Scale bars, 100 μm. (b) Confocal images of colonic myenteric plexus from mice described in (a) at week 1, stained with CC7 (apoptotic cells) and ANNA-1 (neuronal somas) antibodies (supplementary to Fig. 4h). Scale bars, 100 μm.

**Figure S7.**
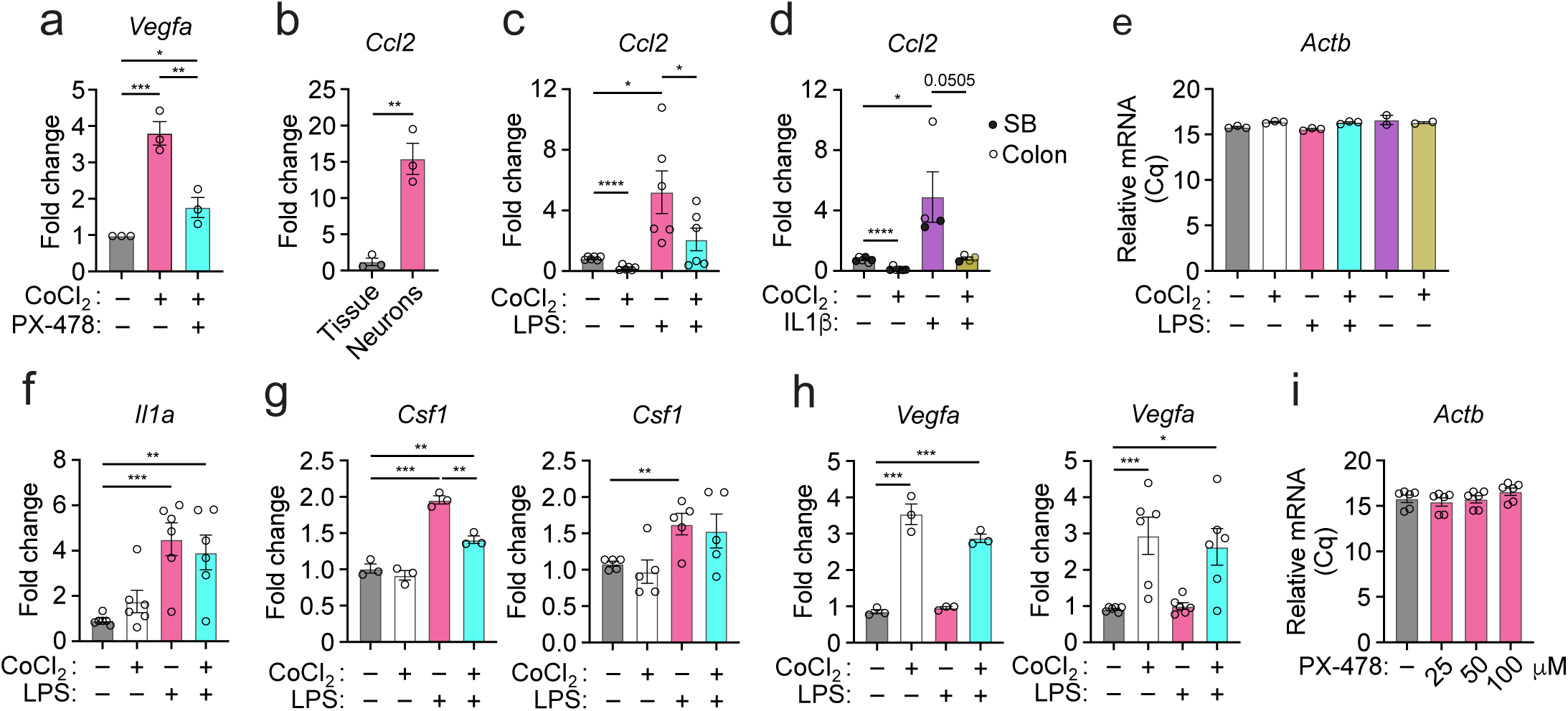
HIF1 signaling in cultured myenteric neurons negatively regulates CCL2 expression. (a) *Vegfa* expression by qPCR in cultured colonic myenteric neurons, treated with CoCl_2_, HIF1α inhibitor PX-478 or their combinations. Each data point is a replica in the same cell culture. (b) *Ccl2* expression by qPCR in cultured embryonic enteric neurons as compared to total adult small bowel muscularis tissue. Each data point is a replica in the same cell culture. (c) *Ccl2* expression by qPCR in cultured colonic myenteric neurons treated with CoCl_2_, LPS or their combination. Each data point is an independent cell culture (supplementary to Fig. 5d). (d) *Ccl2* expression by qPCR in cultured small bowel or colonic myenteric neurons treated with CoCl_2_, IL1β or their combination. Each data point is a replica in the same cell culture. (e) *Actb* mRNA expression (Cq) by qPCR in cultured colonic myenteric neurons treated with CoCl_2_, LPS, IL1β or their combination. Each data point is an independent cell culture. (f) *Il1a* expression by qPCR in the experiment described in (c). Each data point is an independent cell culture (supplementary to Fig. 5f). (g) *Csf1* expression by qPCR in the experiment described in (c). Each data point is a replica in the same cell culture (left) or independent cell culture (right). (h) *Vegfa* expression by qPCR in the experiment described in (c). Each data point is a replica in the same cell culture (left) or independent cell culture (right). (i) *Actb* mRNA expression (Cq) by qPCR in cultured colonic myenteric neurons treated with increasing concentrations of PX478 as indicated. Each data point is a replica in the same cell culture. All graphs show mean ± SEM from 1 representative experiment. Statistical analyses: Unpaired or paired (for pooled independent neuronal cell culture experiments) Student’s t-test, *P≤0.05, **P≤0.01, ***P≤0.001, and ****P≤0.0001.

**Figure S8.**
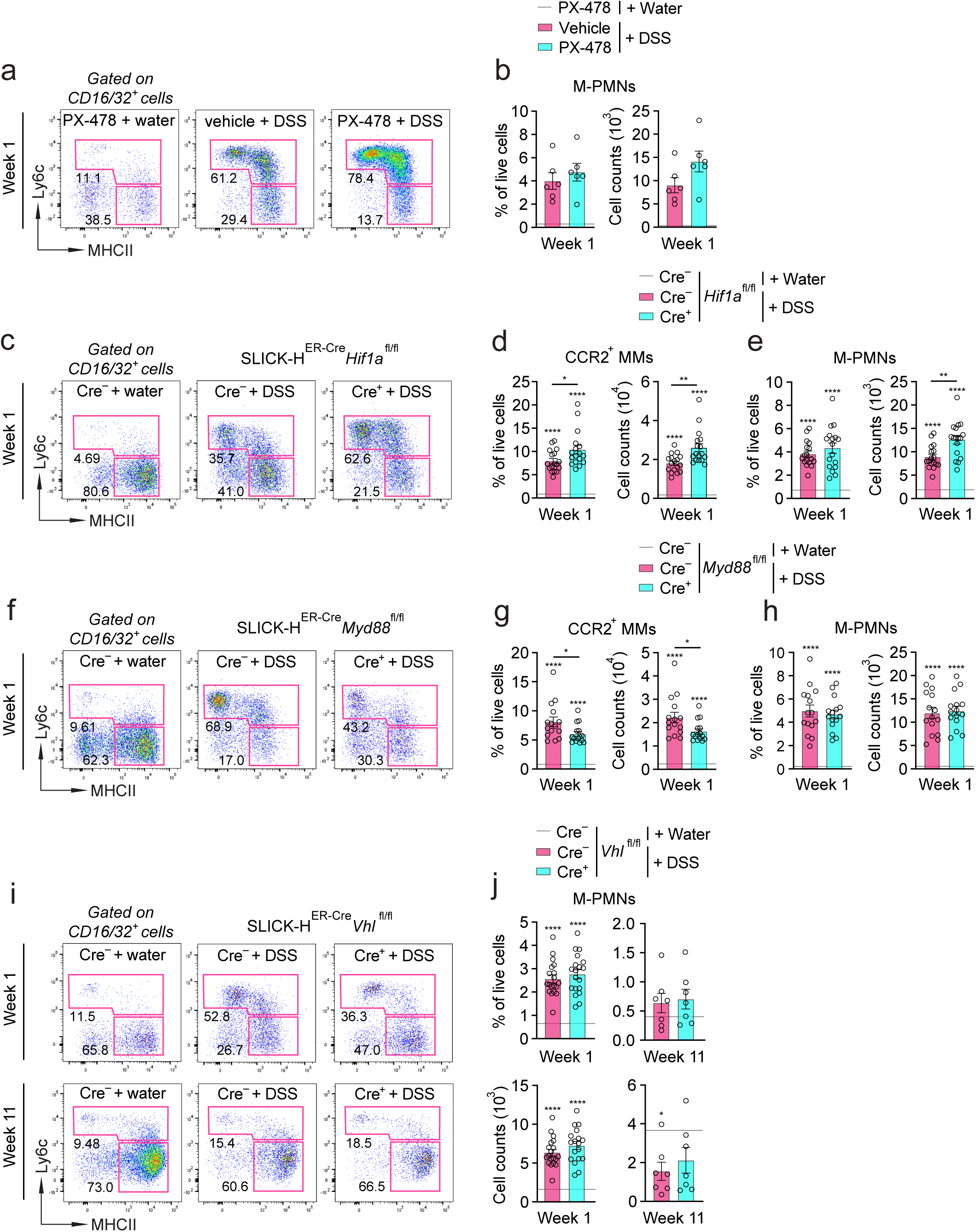
Myenteric inflammation in mice treated with HIF1α inhibitor PX-478 or with neuron-specific depletion of *Myd88*, *Hif1a* or *Vhl*. (a) FC plots showing colonic MM subsets and their percentages in 1×DSS- or water-treated WT mice daily injected with PX-478 or vehicle. (b) Percentage of total viable cells and absolute cell counts of colonic muscularis PMNs (M-PMNs) in mice described in (a), quantified by FC. Baseline shows average value for PX-478-injected mice on water (n=1). (c) FC plots showing colonic MM subsets and their percentages in tamoxifen- and 1×DSS- or water-treated SLICK-H^ER-Cre^*Hif1a*^fl/fl^ (Cre^+^) or *Hif1a*^fl/fl^ (Cre^−^) littermates. (d) Percentage of total viable cells and absolute cell counts of colonic CCR2^+^ MMs in mice described in (c), quantified by FC. Baseline shows average value for tamoxifen- and water-treated *Hif1a*^fl/fl^ littermates (n=13). (e) Percentage of total viable cells and absolute cell counts of colonic M-PMNs in mice described in (c), quantified by FC (baseline n=13). (f) FC plots of colonic MM subsets and their percentages in tamoxifen- and 1×DSS- or water-treated SLICK-H^ER-Cre^*Myd88*^fl/fl^ (Cre^+^) or *Myd88*^fl/fl^ (Cre^−^) littermates. (g) Percentage of total viable cells and absolute cell counts of colonic CCR2^+^ MMs in mice described in (f), quantified by FC. Baseline shows average value for tamoxifen- and water-treated *Myd88*^fl/fl^ littermates (n=12). (h) Percentage of total viable cells and absolute cell counts of colonic M-PMNs in mice described in (f), quantified by FC (baseline n=12). (i) FC plots of colonic MM subsets and their percentages in tamoxifen- and 3×DSS- or water-treated SLICK-H^ER-Cre^*Vhl*^fl/fl^ (Cre^+^) or *Vhl*^fl/fl^ (Cre^−^) littermates (supplementary to Fig. 5j). (j) Percentage of total viable cells and absolute cell counts of colonic M-PMNs in mice described in (i), quantified by FC. Baseline shows average value for tamoxifen- and water-treated *Vhl*^fl/fl^ littermates (baseline n=9 and 5). All graphs show mean ± SEM from 1 representative experiment. Statistical analyses: Unpaired Student’s t-test and two-way ANOVA for multiple comparisons, *P≤0.05, **P≤0.01, ***P≤0.001, and ****P≤0.0001.

**Figure S9.**
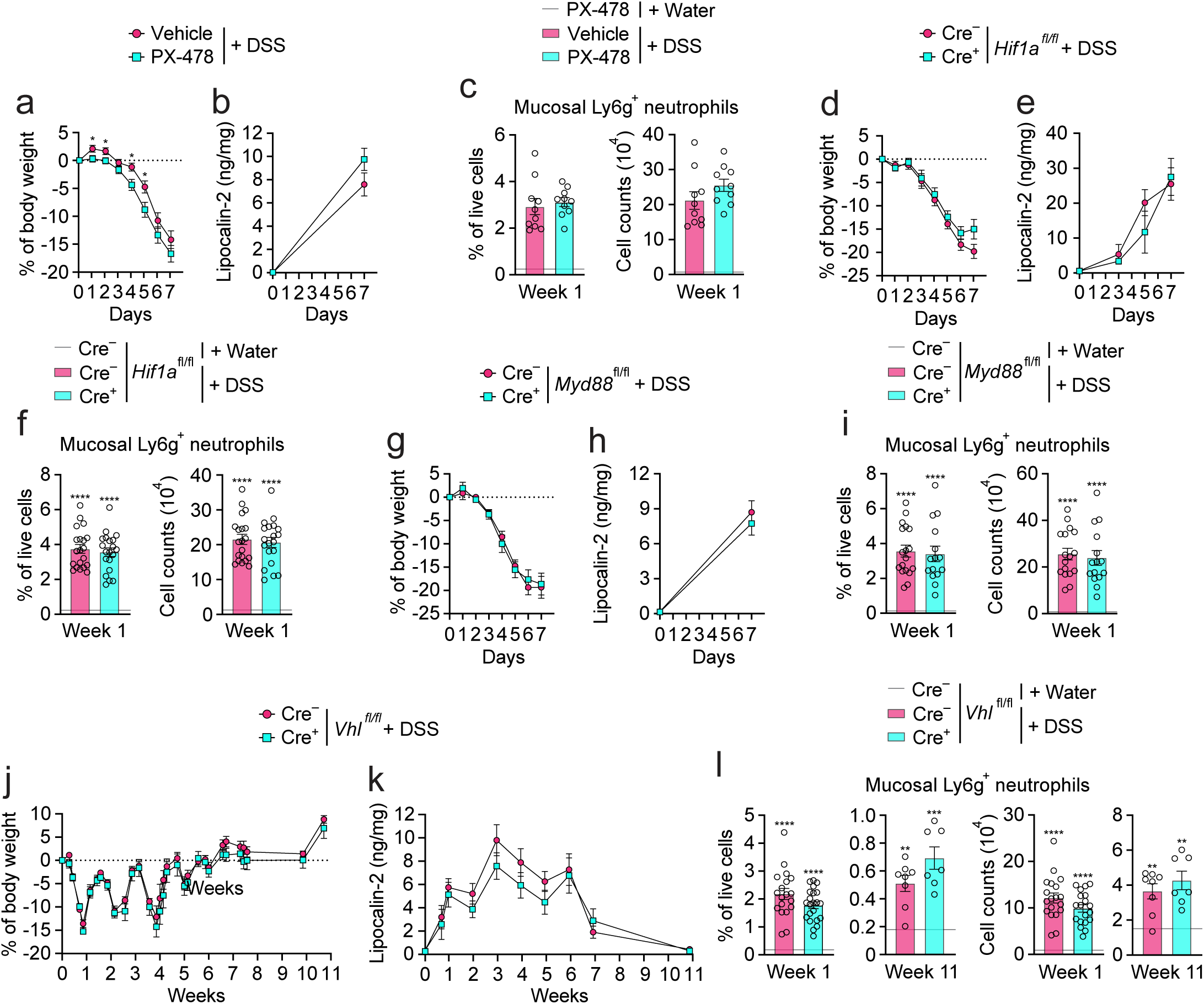
Mucosal inflammation in mice treated with HIF1α inhibitor PX-478 or with neuron-specific depletion of *Myd88*, *Hif1a* or *Vhl*. (a) Percentage of body weight change in PX-478- or vehicle-injected and 1×DSS-treated WT mice. (b) Fecal lipocalin-2 in mice described in (a). (c) Percentage of total viable cells and absolute cell counts of mucosal Ly6g^+^ neutrophils in colons of mice described in (a), quantified by FC. Baseline shows average value for PX-478-injected mice on water (n=1). (d) Percentage of body weight change in tamoxifen- and 1×DSS-treated SLICK-H^ER-Cre^*Hif1a*^fl/fl^ (Cre^+^) or *Hif1a*^fl/fl^ (Cre^−^) littermates. (e) Fecal lipocalin-2 in mice described in (d). (f) Percentage of total viable cells and absolute cell counts of mucosal Ly6g^+^ neutrophils in colons of mice described in (d), quantified by FC. Baseline shows average value for tamoxifen- and water-treated *Hif1a*^fl/fl^ littermates (n=13). (g) Percentage of body weight change in tamoxifen- and 1×DSS-treated SLICK-H^ER-Cre^*Myd88*^fl/fl^ (Cre^+^) and *Myd88*^fl/fl^ (Cre^−^) littermates. (h) Fecal lipocalin-2 levels in mice described in (g). (i) Percentage of total viable cells and absolute cell counts of mucosal Ly6g^+^ neutrophils in colons of mice described in (g), quantified by FC. Baseline shows average value for tamoxifen- and water-treated *Myd88*^fl/fl^ littermates (n=12). (j) Percentage of body weight change in tamoxifen- and 3×DSS-treated SLICK-H^ER-Cre^*Vhl*^fl/fl^ (Cre^+^) and *Vhl*^fl/fl^ (Cre^−^) littermates. (k) Fecal lipocalin-2 in mice described in (j). (l) Percentage of total viable cells and absolute cell counts of mucosal Ly6g^+^ neutrophils in colons of mice described in (j), quantified by FC. Baseline shows average value for tamoxifen- and water-treated *Vhl*^fl/fl^ littermates (n=9 and 5). All graphs show mean ± SEM from 1 representative experiment. Statistical analyses: Unpaired Student’s t-test and two-way ANOVA for multiple comparisons, *P≤0.05, **P≤0.01, ***P≤0.001, and ****P≤0.0001.

**Figure S10.**
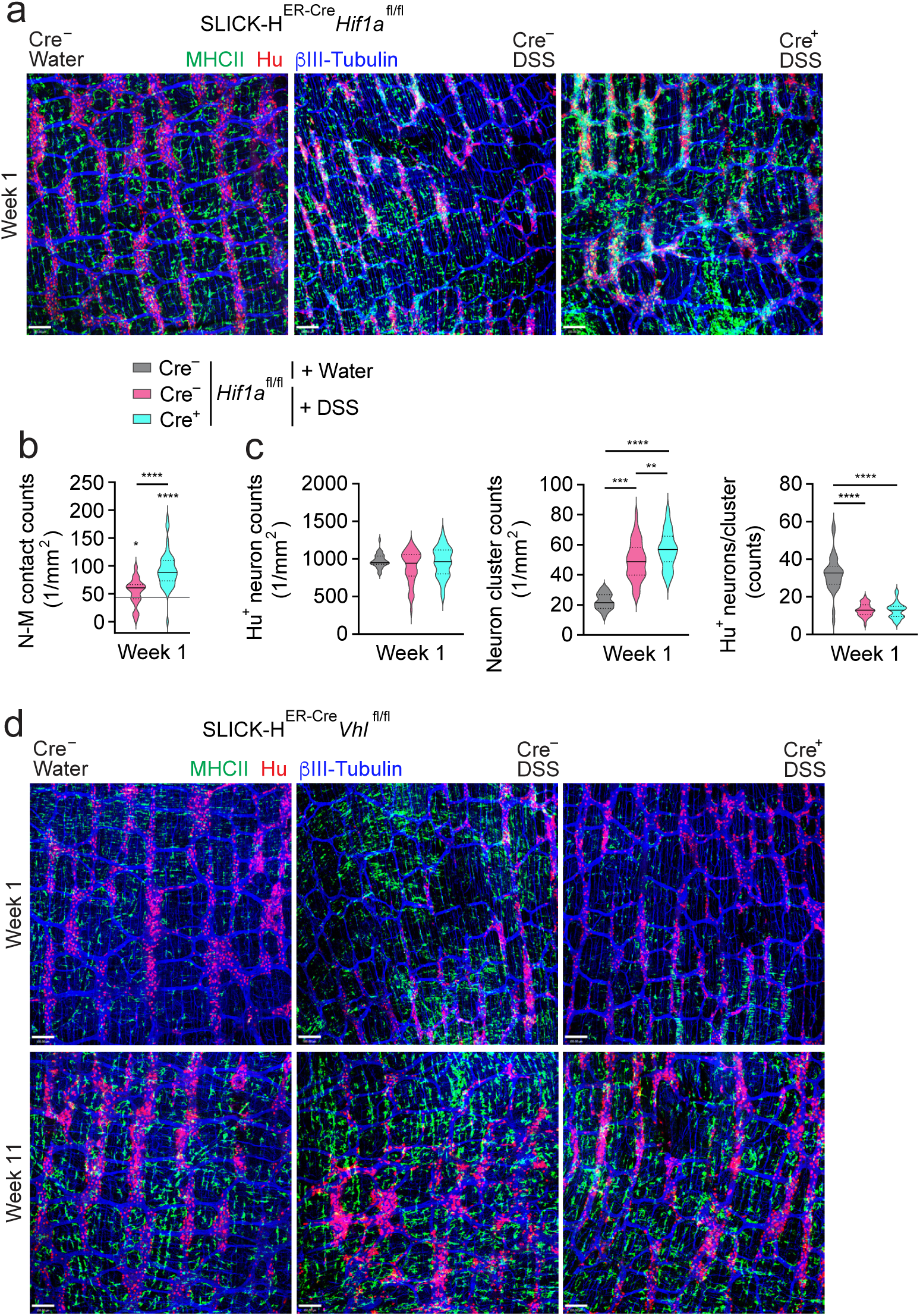
ENS remodeling in mice with neuron-specific depletion *Hif1a* or *Vhl*. (a) Confocal images of colonic myenteric plexus from tamoxifen- and 1×DSS- or water-treated SLICK-H^ER-*Cre*^*Hif1a*^fl/fl^ (Cre^+^) and *Hif1a*^fl/fl^ (Cre^−^) littermates, stained with MHCII (MMs), Hu (neuronal somas) and ýIII-Tubulin (nerve fibers) antibodies. Scale bars, 100 μm. (b) Structural changes in myenteric plexus of mice described in (a) quantified by confocal microscopy. Data shown as a distribution of pooled values per group. (c) Quantification of contacts between MMs and myenteric neurons in mice described in (a) by confocal microscopy. Baseline shows average value for tamoxifen- and water-treated *Hif1a*^fl/fl^ littermates (n=6). (d) Confocal images of colonic myenteric plexus from tamoxifen- and 3×DSS- or water-treated SLICK-H^ER-*Cre*^*Vhl*^fl/fl^ (Cre^+^) and *Vhl*^fl/fl^ (Cre^−^) littermates, stained with MHCII (MMs), Hu (neuronal somas) and ýIII-Tubulin (nerve fibers) antibodies. Scale bars, 100 μm.

## METHODS

### Experimental animals

Wild type (WT) C57BL/6 (B6) mice were obtained from Charles River Laboratories. *Il10*^−/−^ mice (B6.129P2-*Il10^tm1Cgn^*/J; Jackson Laboratory [JAX] #002251)^1^, *Nestin*^ER-Cre^ (C57BL/6-Tg(Nes-cre/ERT2)KEisc/J; JAX #016261)^2^, *Rosa26*-STOP^fl/fl^tdTomato (B6.Cg-*Gt(ROSA)26Sor^tm^*^14^*^(CAG-tdTomato)Hze^*/J; JAX #007914)^3^, *Ccr2*^RFP/RFP^*Cx3cr1*^GFP/GFP^ (B6.129(Cg)-*_Cx3cr1tm1Litt Ccr2tm2.1Ifc_*_/JernJ; JAX #032127)4,5, *Ccr2*ER-Cre/ER-Cre (C57BL/6-*Ccr2*_*_em1(icre/ERT2)Peng_*_/J;_ JAX #035229)^6^, SLICK-H^ER-Cre^ (Tg(Thy1-cre/ERT2,-EYFP)HGfng/PyngJ; JAX #012708)^7^, *Ccl2*-RFP^fl/fl^ (B6.Cg-*Ccl2^tm^*^1^*^.1Pame^*/J; JAX #016849)^8^, *Myd88*^fl/fl^ (B6.129P2(SJL)-*Myd88^tm1Defr^*/J; JAX #008888)^9^, *Hif1a*^fl/fl^ (B6.129-*Hif1a^tm3Rsjo^*/J; JAX #00756)^10^ and *Vhl*^fl/fl^ (C;129S-*Vhl^tm1Jae^*/J; JAX #004081)^11^ mice were purchased from the Jackson Laboratory. Since male mice were shown to be more susceptible to DSS-induced colitis^12^, only adult females between 5-8 weeks of age were used. To minimize the role of genetic background, and reduce cage effects and differences in microbiota, cohoused littermates were used. Mice were maintained in specific pathogen-free conditions with food and water provided ad libitum. All animal experiments were performed in accordance with and approved by the Institutional Animal Care and Use Committees (IACUC) at Penn State University College of Medicine, Hershey, PA, the University of Massachusetts Chan Medical School, Worcester, MA, and the Johns Hopkins University School of Medicine, Baltimore, MD.

### Colitis induction

To induce colitis, mice received 2-4% dextran sodium sulfate (DSS) in sterile drinking water for 5 days and then switched to sterile drinking water for 5 days. This constituted 1 cycle of DSS treatment (1×DSS). For most experiments, mice were given 3 cycles of DSS (3×DSS) followed for up to 60-80 days of post-colitis recovery period. Control mice received sterile drinking water throughout the experiment. For analysis, mice were euthanized on days 5-8 for an early acute colitis phase and days 60-80 for the recovery phase after starting the DSS treatment regimen. For spontaneous colitis, mice received sterile drinking water from the weaning stage and were monitored weekly for body weight and clinical symptoms until they presented with rectal prolapse indicating progressive colitis.

### Inducible gene depletion and cell fate mapping *in vivo*

Inducible Cre mice and corresponding controls were orally gavaged with 20 mg/ml of tamoxifen (Sigma T5648-5G) dissolved in corn oil to induce Cre recombinase expression. Before starting the 1×DSS or 3×DSS treatment regimens, mice received two 5-day cycles of tamoxifen with a 5-day break between each cycle. After starting the DSS treatment, mice were given 2 consecutive gavages every 10 days until the end of the experiment. To trace monocyte-derived MMs, tamoxifen was given for 4 days before the start of the DSS treatment and then given on alternate days until analysis.

### **HIF1α** inhibition *in vivo*

To inhibit HIF1α *in vivo*, B6 mice were injected with PX-478 (#202350, MedKoo Biosciences) in sterile 1×PBS intraperitoneally (i.p.) daily for 3 days before the start of DSS treatment at a dosage of 25 mg/kg and then continued daily throughout the DSS cycle until analysis at week 1^13,14^. Control mice were given i.p. injections of sterile 1×PBS daily until analysis.

### Detection of tissue hypoxia

To detect hypoxic cells *in vivo* during colitis, the EF5 Hypoxia Detection Kit that included EF5 reagent and EF5-specific antibody (EF5-30A4, EMD Millipore) was used^15^. Briefly, at week 1, 3 hrs before euthanasia, control and DSS-treated mice were retro-orbitally injected with 10 nM of EF5 compound. Harvested colons were washed with complete HBSS (HBSS with 2% FBS) containing dithiothreitol (DTT), rolled into “Swiss rolls”, embedded into Optimal Cutting Temperature (OCT) compound, and snap-frozen at –80°C. Eight μm tissue cross-sections of the frozen “Swiss roll” tissue were fixed in 4% paraformaldehyde (PFA), blocked and stained with the antibody specific for Hu (1:500, clone EPR19098, Abcam, stains neuronal somas) followed by donkey anti-rabbit Cy3 secondary antibody (1:800, Jackson Immunoresearch) and DAPI. Tissue sections were re-fixed with 4% PFA, blocked, stained with Alexa Fluor 488 conjugated EF5 antibody (clone ELK3-51, 150 μg/ml) and mounted onto histological slides with Prolong Diamond Antifade mounting medium (#P36965, Thermo Fisher).

### Sample and tissue collection

To monitor luminal lipocalin-2 levels, fecal pellets were collected weekly and at the end of the experiment, resuspended in 1×PBS and stored at –80°C for later analysis. Blood was drawn by cardiac puncture to analyze blood monocyte populations by flow cytometry. Colons were processed as we described previously^16,17^. Briefly, harvested colons were cleaned and washed using complete HBSS containing DTT and EDTA to remove mucus and epithelial cells. Colonic muscularis externa was mechanically separated from the mucosa and submucosa as described^16,17^.

### Tissue processing for immunofluorescence

A 5 mm long piece of the muscularis externa from the mid colon was excised and fixed with 4% PFA. Tissues expressing fluorescent proteins from reporter mice (*Ccl2*-RFP^fl/fl^ mice containing mCherry, *Rosa26*-STOP^fl/fl^tdTomato mice containing tdTomato and *Ccr2*^RFP/+^*Cx3cr1*^GFP/+^ mice containing RFP and GFP) were fixed with 4% PFA containing 20% sucrose to preserve the fluorescent proteins. Post-fixation, tissue was stored in 1×PBS containing 30% sucrose at 4°C until immunofluorescent staining.

### Human intestinal tissue

Deidentified full-thickness intestinal specimens surgically removed from 3 patients with CD and 3 non-IBD control patients with colorectal cancer (CRC) undergoing colectomy were provided by the UMass Center for Clinical and Translational Science (UMCCTS) Biorepository under an approved IRB protocol. For CD, non-inflamed and inflamed regions were provided for each patient. For two CD patients, large bowel tissue from non-inflamed and inflamed regions defined macroscopically by a Board-certified pathologist were provided. For one CD patient, the uninflamed sample was from colon, while the inflamed sample was from the terminal ileum; the terminal ileum sample was excluded from further analysis. For CRC patients, normal large bowel regions adjacent to tumors were used as a control.

Received tissue samples were rinsed with 1×PBS, fixed with 4% PFA and incubated in 1% PFA with increasing concentrations (10%, 20% and 30%) of sucrose. Next, tissues were incubated in a 1:1 mixture of OCT and 1% PFA containing 30% sucrose and embedded in OCT compound before freezing at –80°C.

### Immunofluorescence

#### Mouse whole tissue

Fixed 5 mm piece of intact muscularis externa was washed with 1×PBS, blocked in 1×PBS containing 2% BSA and 1% Triton X, stained with a specific primary antibody cocktail, washed again, and stained with a mixture containing DAPI and a combination of fluorochrome-conjugated antibodies raised in donkey against rat, goat, rabbit, chicken, human or Armenian hamster. Stained tissues were mounted on histological slides with Prolong Diamond Antifade mounting medium. To stain cells of the myenteric plexus, antibodies specific to mouse Hu (stains neuronal somas, 1:500, clone EPR19098, Abcam), βIII-Tubulin (stains nerve fibers, 1:250, rabbit or chicken polyclonal, Abcam), PGP9.5 (stains entire neurons, 1:500, clone EPR4118, Abcam), GFAP (stains glial cells, 1:1000, chicken polyclonal, Abcam), CD31 (stains endothelial cells, 1:200, clone MEC7.46, Novus Biologicals) and MHC Class II (MHCII) I-A/I-E (stains MMs, 1:100, clone MS/114.15.2, Biolegend) were used. Serum collected from a seropositive patient with a confirmed high titer of type-1 antineuronal nuclear antibody (ANNA-1) specific to Hu (stains neuronal somas, 1:1000, reactive to mouse and human) was provided by Dr. Sean J. Pittock, at Mayo Clinic. Antibodies specific for tdTomato, RFP or mCherry (1:250, goat polyclonal, Origene) and for GFP (1:100, clone FM264G, Biolegend) were used to amplify the signal of endogenously expressed fluorescent reporter proteins. To stain apoptotic neurons, fixed colonic muscularis externa was blocked in 1×PBS containing 5% donkey serum, 0.3% Triton X and 1% penicillin-streptomycin, then stained with primary antibodies specific for cleaved caspase 7^18^ (CC7, 1:250, clone Asp 198, Cell Signaling), βIII-Tubulin and ANNA-1 followed by staining with fluorochrome-conjugated secondary antibodies and mounting as previously described. To stain CCL2-expressing cells, 0.25 mg of Brefeldin A (Enzo Life Sciences, Farmingdale, NY) was i.p. injected into DSS-treated mice 6 hrs before euthanasia. Colons were harvested, processed, and fixed as described above. Ten μg/ml of Brefeldin A was maintained in all buffers until the blocking step^19^. Fixed tissues were blocked, stained with a primary antibody cocktail containing antibodies specific for RFP, Hu and MHCII followed by washing, staining with fluorochrome-conjugated secondary antibodies, and mounting on histological slides.

#### Human tissue sections

Ten μm thick intestinal tissue cross-sections were prepared and blocked with 5% donkey serum in 1×PBS containing 0.05% Triton X. Sections were stained with primary antibodies specific for mouse Hu (1:250), βIII-Tubulin (1:250) and human HLA-DR (1:250, LN-3, Leica/Novocastra) followed by staining with a cocktail of DAPI and fluorochrome-conjugated secondary antibodies and mounting on histological slides.

### Microscopy

Whole muscularis externa or cross-sections of mouse colons were imaged using the Leica HC PL APO CS2 10×/0.4 NA objective at 1.136 μm x1.136 μm pixel size or 0.568 μm x 0.568 μm pixel size on the Leica SP8 confocal micropscope. The Leica HC PL APO CS2 20×/0.75 NA objective was used at 0.284 μm x 0.284 μm pixel size on the Leica SP8 confocal micropscope. For 3D imaging of neuron-macrophage interactions, the Nikon CFI75 Apo LWD 25x W 1.1 NA objective on the Nikon A1R multiphoton was used. The 10× and 20× water immersion and 63× oil immersion objectives on the Nikon A1R multiphoton microscope were used. All images were processed using Volocity 6.3.1, Quorum Technologies Inc. Human tissue sections were imaged using a 20× 0.75 NA Zeiss Plan Apo objective on the TissueGnostics TissueFAXS SL Q slide scanning microscope. Images were acquired in fluorescence mode and focus points across the selected region were determined using the DAPI channel. Acquired images were processed using StrataQuest, TissueGnostics, Inc.

### Image analysis to quantify ENS remodeling

#### Confocal microscopy of mouse myenteric plexus

ENS remodeling was assessed by automated analysis of XY-images of intact myenteric plexus from mouse colons generated by confocal microscopy to quantify the (i) total number of Hu^+^ neurons, (ii) number of Hu^+^ neuron clusters, (iii) number of Hu^+^ neurons per cluster, (iv) number of interganglionic fiber tract (IGFT) regions, (v) IGFT region size, (vi) number of contacts between MMs and neurons and (vii) total MHCII^+^ MM area per field of view (FOV) using Volocity 6.3.1, Quorum Technologies Inc. All data were normalized to mm^2^. For all experiments, 5-8 FOVs were analyzed per mouse and each experiment had n ≥ 3 mice per group.

Neuron and ganglia identification criteria were adapted from the methodology previously described^20–23^. Briefly, an intact *ganglion* or a smaller *cluster of neurons* was defined as a contiguous group of 2 or more Hu ^+^ neurons separated from other Hu^+^ neurons by a distance ≥ the diameter of at least 2 neurons. The neuronal ganglion or cluster must also be surrounded by ýIII-Tubulin^+^ nerve fibers. *Neuron counts per cluster* were defined as the number of neurons identified in a ganglion or cluster. MMs were identified as MHCII^+^ cells and were quantified by measuring the total signal intensity area (i.e., *total MHCII^+^ area*). Colocalization of MHCII^+^ signal with Hu^+^ signal was defined as *contacts* and was measured by identifying the MHCII signal and colocalizing it with the Hu^+^ signal and conversely. Both ways of quantification were found to be similar and comparable. We quantified the organization of nerve fiber architecture of the myenteric plexus by counting the number and measuring the area of the rounded spaces between myenteric ganglia and ýIII-Tubulin^+^ IGFT that we referred to as *IGFT regions*. A combination of all readouts was the most informative, with the total number of neurons being the least sensitive and requiring a higher sample size

#### Multiphoton microscopy of mouse myenteric plexus

Automated analysis of z-stack multiphoton images was done to quantify the (i) number of Hu^+^ neurons, (ii) number of MHCII^+^ MMs, (iii) number of contacts between MHCII^+^ MMs and Hu^+^ neurons and (vii) Hu^+^ neuronal signal inside MHCII^+^ MMs that measured uptake of neuronal cellular components by MMs, per FOV using Volocity 6.3.1, Quorum Technologies Inc. All data were normalized to mm^2^. For all experiments, 5-8 FOVs were analyzed per mouse and each experiment had n ≥ 3 mice per group.

#### ENS remodeling in human large bowel cross-sections

Analysis of immunofluorescence images was performed using a StrataQuest v7.1.1.143 analysis pipeline that was developed to quantify the (i) total area of HLA-DR^+^ signal per region of interest (ROI) indicating inflammation, (ii) total area of βIII-Tubulin^+^ signal per ROI indicating innervation, and (iii) area of Hu^+^ neuronal ganglia. DAPI channel was used to identify nuclei. Hu^+^ neurons were manually counted within each ganglion to calculate the density of Hu^+^ neurons per ganglia area (*neuron density*). All data were normalized to mm^2^.

### Flow cytometry and cell sorting

Freshly isolated mouse colonic muscularis externa or mucosa with submucosa were subjected to enzymatic digestion to obtain single-cell suspensions as we described^16,17^. The number of total live cells in suspension was calculated using a hemocytometer. These total cell counts were used to calculate absolute counts of different subsets. Blood samples were subjected to red blood cell lysis. Obtained single-cell suspensions were stained with antibodies for cell-specific markers.

#### Antibodies

All antibodies were purchased from Biolegend (San Diego, CA) unless indicated. To stain neurons, antibodies specific for mouse CD24 (Clone M1/69, BD Biosciences) and CD90.2 (Clone 53.-2.1, BD Biosciences) were used. To stain monocytes and macrophages, antibodies specific to mouse CD45 (clone 30-F11), CD11b (clone M1/70), CD11c (clone N418), CD16/32 (Clone 93), Ly6c (clone HK1.4) and MHCII I-A/I-E (clone MS/114.15.2) were used. To stain neutrophils an antibody specific for mouse Ly6g (clone 1A8) was used. To stain B cells and T cells, antibodies specific for mouse B220 (clone RA3-6B2) and CD3 (Clone 500.A2, Thermo Fisher) were used. For live cell staining either DAPI (Biolegend) or Aqua fluorescent reactive dye (Invitrogen) were used. For intracellular staining of PGP9.5 expressed by enteric neurons, fresh cells were first stained for cell surface markers, fixed using the BD Cytofix/Cytoperm™ Fixation/Permeabilization Solution Kit and then stained with antibody specific to mouse PGP9.5. All stained samples were acquired on BD LSRII™ Flow Cytometers.

#### Gating strategy

In the mucosa with submucosa, immune cell subsets were analyzed using a published gating strategy^16,17,19^. In the muscularis, CD90^+^CD24^+^ neurons and all other non-neuronal stromal cells (CD90^+^, CD24^+^, CD90^−^CD24^−^subsets) were defined by gating on CD45^−^ live singlets. Immune cell subsets were defined by gating on CD45^+^ live singlets. Total CD45^+^ lymphocytes were defined as CD45^+^CD11b^−^ cells. Muscularis T cells were defined as CD45^+^CD11b^−^CD3^+^ cells. CD45^+^CD11b^+^ myeloid cells were divided into muscularis polymorphonuclear (M-PMNs) cells defined as SSC-A^hi^CD16/32^−^ (Ly6g^−^ and Ly6g^+^) cells and MMs defined as SSC-A^lo-int^Ly6g^−^CD16/32^+^ cells. MMs were further divided into additional subsets based on the combination of *Ccr2*-RFP, *Cx3cr1*-GFP and cell surface binding of Ly6c and MHCII antibodies as shown in the Results.

#### Flow cytometry analysis

Data analysis was performed using FlowJo software (FlowJo LLC, Ashland, OR). *Ccr2*^RFP/+^*Cx3cr1*^GFP/+^ reporter mice were used to perform an unbiased comparative analysis of immune cell subsets of the muscularis externa. All CD45^+^ events were concatenated group-wise. Unsupervised t-SNE analysis was performed with FlowJo t-SNE plugin set to 6 phenotypic markers (CD11b, CD16/32, *Cx3cr1*-GFP, *Ccr2*-RFP, Ly6c, MHCII) and used the following settings: Auto (opt-SNE) learning algorithm with approximate (random projection forest-ANNOY) KNN algorithm^24^ and FFT Interpolation (flt-SNE) gradient algorithm.

#### Sorting of MMs

*Ccr2*^RFP/+^*Cx3cr1*^GFP/+^ reporter mice were used to sort MM subsets for bulk RNA sequencing. Colonic muscularis externa was isolated from cohorts of 3-5 DSS-treated and 3-5 water-treated mice at week 1 time point. Muscularis single-cell suspensions from 5 mice were pooled per condition and represented one replicate. Specific MM subsets were isolated by FACS.

In total, 5 such replicates were prepared representing 5 independent cell sorting experiments. Cell sorting was done as we described previously^16,17^. Cells were stained with a combination of CD45, CD11b, CD16/32, Ly6c, MHCII antibodies. MMs were defined as CD45^+^CD11b^+^CD16/32^+^ viable singlets, and MM subsets were sorted by a 4-way sort using the BD FACSAria™ Fusion flow cytometer. Based on *Ccr2*-RFP expression, MM subsets were broken down into CCR2^+^MHCII^−^ (subset 1), CCR2^+^Ly6c^+^MHCII^+^ (subset 2), CCR2^+^Ly6c^−^MHCII^+^ (subset 3), CCR2^−^Ly6c^−^ MHCII+ (subset 4) and CCR2^−^MHCII^−^ (subset 5) subsets. Between 5,000 and 10,000 cells of subsets 2, 3, 4, 5 were sorted from control mice, and subsets 1, 2, 3, 4 were sorted from DSS mice, based on cell frequency.

#### Sorting of neurons

Single-cell suspensions were prepared from the muscularis of WT B6 mice with or without colitis. Non-immune cells were defined as CD45^−^ viable singlets. CD90^+^CD24^+^ neurons and other CD45^−^ cells (pooled CD24^−^CD90^+^, CD90^−^CD24^+^, CD90^−^CD24^−^ stromal cell subsets) were subjected to a 4-way sort using the BD FACSAria™ Fusion flow cytometer. Sorted cells were collected in RPMI with 10% FBS and stored in TRIzol at –80°C for further processing and analysis.

### RNA isolation and gene expression analysis

Total RNA was extracted from sorted cells using TRIzol extraction protocol as described previously^19^. RNA was reverse transcribed using RNA to cDNA Ecodry premix (Takara Bio, Kusatsu, Japan). Gene expression was measured using Power SYBR Green PCR Master mix (Life Technologies) on a BIO-RAD qPCR instrument. *Ccl2* (Mm.PT.58.42151692), *Vegfa* (Mm.PT.58.13368357), *Csf1* (Mm.PT.58.11661276), *Il1a* (Mm.PT.58.32778767), *Hif1a* (Mm.PT.58.12608714), *Elavl3* (Mm.PT.58.30335452), *Elavl4* (Mm.PT.58.33020194) and *Actb* (IDT 419680055, 419680054) PrimeTime qPCR primers were purchased from Integrated DNA Technologies (Skokie, IL). Gene expression levels were calculated by normalization to *Actb* and shown as fold change over the control group or population described in the figure legends. C_q_ values for *Actb* for each cell culture condition were used as an indirect measurement of cell viability in culture.

### mRNA sequencing

Total RNA was isolated from sorted cell populations as described previously. cDNA libraries were prepared using the QuantSeq 3’mRNASeq Library Prep Kit FWD for Illumina (Lexogen, Vienna, Austria) following the manufacturer’s instructions. Briefly, RNAseq libraries were prepared using Lexogen’s QuantSeq 3’ mRNASeq V2 Library Prep Kit Forward (FWD) with 12 nucleotide Unique Dual Indices (UDIs) following the manufacturer’s instructions for low input RNA library prep. One nanogram of total RNA was reverse-transcribed using oligo (dT) primers. The second cDNA strand was synthesized by random priming followed by cDNA purification and library amplification using Lexogen’s 12 nt Unique Dual Indices and 24 cycles. The libraries were analyzed for size distribution and concentration using BioAnalyzer High Sensitivity DNA kit (Agilent Technologies, Santa Clara, CA). Libraries were pooled at equimolar concentrations and sequenced on Novaseq6000 (Illumina) to get approximately 5 million paired end 50 bp reads.

### mRNAseq analysis of sorted MM subsets

To analyze data generated from mRNAseq of sorted MM subsets, FastQC (v0.11.9) was used to compute basic statistics and check quality control (QC) of paired reads from RNAseq^35^. The results showed that R2 of the paired reads have overall low quality, and considerable number of overrepresented repetitive sequences, thus, only R1 reads were used for further analysis. Trimmomatic (v0.36) was used to trim low-quality parts of reads^25^. STAR (v2.7.5b) was used to align reads to GRCm39^26^. Reads counts for each gene were summarized using RNA-SeQC (v2.1.0)^27^. Contribution of covariates to the variance were investigated using variancePartition (v1.32.2)^28^. The differentially expressed genes were identified by limma-voom model implemented in edgeR package (v4.0.14)^29^. The signature for each macrophage subset was generated by taking the union of all differential expression signatures (including up- and down-regulated) from pairwise comparison of each subset to all other subsets. Additional differential expression signatures generated include DSS versus wild-type control for each subset and comparisons of specific pairwise DSS subgroups. The cutoff for significance is less than the adjusted *p* value 0.05 and greater than log fold change 1.5 or less than –1.5. The heatmap was generated using heatmaply (v1.5.0)^30^. Pathway analyses were conducted using EnrichR (v3.2)^31^ and David (https://david.ncifcrf.gov/home.jsp)^31^.

### Single-cell RNA sequencing of muscularis externa

Single-cell suspensions were made from the muscularis externa of the colonic muscularis externa isolated from 11-week-old adult WT B6 mice as described previously^20,23^. The separated muscularis externa was digested, filtered, and resuspended into maintenance solution containing OptiMEM with GlutaMax, B27 supplement, Actinomycin D and RNAse inhibitor. Filtered single-cell suspensions were assessed for viability and cell counts using Trypan Blue and Cell Countess (Invitrogen). The library was prepared using Chromium Next GEM Single Cell 3’ kit v3.1 (10x genomics) at a read depth of 20,000/cell and was sequenced using the Illumina NovaSeq 6000 by the Johns Hopkins Single Cell Sequencing Core.

### Transcriptome analysis of muscularis externa by single-cell RNA sequencing

To analyze data generated from the scRNAseq of muscularis externa, raw 10× read processing and QC raw sequence reads were quality-checked using FastQC (v0.11.9) software^48^. The Cell Ranger version 7.0 software suite from 10x Genomics was used to process, align, and summarize unique molecular identifier (UMI) counts against the mouse GRCm38 assembly reference genome analysis set, obtained from https://www.10xgenomics.com. Filtered count matrices from Cell Ranger were imported into R (version 4.3.2) for further processing. Low quality cells were filtered, such as cells for which a high percentage of UMIs originated from mitochondrial features (more than 20%) and cells with fewer than 200 expressed genes. Three methods: Doubletfinder (v2.0.4)^32^, Scds (v1.18.0)^33^ and scDblFinder(v1.16.0)^34^ were applied to detect doublets, and doublets cells identified by at least two methods were filtered. Median absolute deviation method was used to derive a cutoff for the maximum number of genes in cell. After filtering, 34,417 cells were left, and used for further analysis. The Seurat R package (version 5.0) was used for normalization. Harmony (v1.2) was used for integration and plotting^35,36^. Slingshot (v2.10) was used for trajectory and pseudotime analysis^37^. CellChat (v2.1.2) was used to predict cell-to-cell communication between macrophage clusters and neurons^38^.

### Human bulk RNA sequencing analysis

#### Bayesian network analysis

We curated a hypoxia gene list to represent a signature of genes involved in hypoxia signaling combined from^39^ and Enrichr^31,40,41^. To identify genes expressed in inflamed enteric neurons, we developed a method to generate cell type specific signatures as described in Lu et al, (manuscript in preparation), using scRNAseq from the previously published data set^42^. We generated hypoxia subnetworks in the following Bayesian networks (BN): control ileum BN, CD ileum BN, control colon BN, CD colon BN, ulcerative colitis (UC) colon BN and, by projecting each signature onto the BN, identifying all overlapping nodes between the signature and the BN, then extending out two pathlengths, and extracting the largest connected subnetwork using Cytoscape 3.10.2. To test for enrichment of the signature in the sub-network relative to the full BN, we performed the Fisher’s Exact test, using p value of < 0.05 as a cut off.

#### CCL2 subnetwork analysis

The *CCL2* subnetworks are comprised by identifying the *CCL2* node and extending out from *CCL2* to include all directly connected nodes within three path lengths in the following IBD and control Bayesian networks: (i) *CCL2* sub-network of the CD ileum sub-network, (ii) *CCL2* subnetwork of the CD colon sub-network, (iii) *CCL2* sub-network the UC sub-network and (iv) *CCL2* sub-network of the control colon subnetwork. The Bayesian networks were constructed from RNAseq profiling and DNA SNP panels from intestinal biopsies collected in the Mount Sinai Crohn’s and Colitis Registry (MSCCR) cohort at Mount Sinai^43,44^, as previously described^45^. Cytoscape (v3.10.2) was used to generate the *CCL2* subnetwork plots.

### Total gastrointestinal (GI) transit time assay

Total GI transit time assay was performed as we described^46^. Briefly, 3 days prior to the assay, mice were acclimated by being individually housed with hydrogel, no bedding and fasted for the duration of the assay. On the assay day, mice were given 300μl of sterile water containing 6% carmine red (Sigma-Aldrich), 0.5% methyl-cellulose (Sigma-Aldrich) and 0.9% NaCl by intragastric gavage, and fecal pellets were examined for the first appearance of the red dye. Total GI transit time was calculated as the time between gavage and the appearance of the first red fecal pellet.

### Embryonically derived enteric neuron culture

Embryonically derived enteric neuron culture was prepared from intestines isolated from B6 WT embryos on embryonic day 17 as described^46^.

### Primary adult enteric neuron culture

Colonic muscularis externa of 6-week-old mice were extracted, and single-cell suspensions were prepared as described above. Cells were seeded into Matrigel-coated 24-well plates at a density of 150,000 cells/well and maintained in complete neuronal culture media containing Neurobasal-A media (Gibco), L-Glutamine (200mM, Gibco), B27 (1×, Gibco), glial-derived neurotrophic factor (GDNF, 10 ng/ml, Shenandoah), FBS (1%), Amphotericin B (2.5 μg/ml, Gibco) and Penicillin-Streptomycin (1%, Gibco). Morphological changes were observed daily, and on days 13-14 differentiated neurons in culture were used for experiments.

### Chemical hypoxia *in vitro*

Cobalt chloride (CoCl_2_) was used as a chemical mimetic to simulate hypoxia^47^ *in vitro*. Differentiated enteric neuron culture was treated with CoCl_2_ (200μM, Sigma-Aldrich) for 12 hrs followed by treatment with lipopolysaccharide (LPS, 100 ng/ml, Sigma-Aldrich) or recombinant murine IL1β (100 ng/ml, #211-11B, PeproTech) for another 9 hrs. To inhibit HIF1α during chemical hypoxia, differentiated enteric neuron cultures were pretreated with PX-478 (25 μM, #202350, MedKoo Biosciences) for 16 hrs before inducing chemical hypoxia using CoCl_2_ for 12 hrs followed by LPS treatment for 9 hrs. PX-478 was maintained in the culture by replenishing it every 16 hrs. At the end of the experiment, cells and supernatants were collected for gene expression and protein analyses.

### Protein expression by ELISA

The concentration of fecal lipocalin-2 was measured using R&D Systems ELISA kit (#DY1857). The concentration of secreted CCL2 in supernatants of cultured enteric neurons was measured using R&D Systems ELISA kit (#DY479-05).

## ACKNOWLEDGMENTS

We thank the Sanderson Center for Optical Experimentation (SCOPE) (RRID:SCR_022721) and the Flow Cytometry Core (RRID:SCR_012630) at the University of Massachusetts Chan Medical School (UMCMS) for their assistance, the Genomic Sciences Core (RRID: SCR_021123) and the Advanced Light Microscopy Core (RRID:SCR_022526) at the Penn State University College of Medicine (PSUCoM) for their assistance. The Genome Sciences Core (RRID:SCR_021123) services and instruments used in this project were funded, in part, by the Penn State University College of Medicine via the Office of the Vice Dean of Research and Graduate Students and the Pennsylvania Department of Health using Tobacco Settlement Funds (CURE). The content is solely the responsibility of the authors and does not necessarily represent the official views of the University or College of Medicine. The Pennsylvania Department of Health specifically disclaims responsibility for any analyses, interpretations or conclusions. The content is solely the responsibility of the authors and does not necessarily represent the official views of the University or College of Medicine. The Pennsylvania Department of Health specifically disclaims responsibility for any analyses, interpretations or conclusions. The library preparation and scRNA sequencing of colonic cells was performed at the Single Cell and Transcriptomics Core at the Johns Hopkins University.

This work was supported by Junior Faculty Research Scholar Award by PSUCoM and Pennsylvania Department of Health using Tobacco CURE Funds (M.B.), Kenneth Rainin Foundation Innovation Award (M.B.), HIH-NIAID R21AI126351-01 (M.B.), NIH-NIDDK R01DK107603 (M.B.), NIH-NIA R01AG066768 (Su.K.), NIH-NIA R21AG072107 (Su.K), the Harvard Digestive Disease Center Pilot and Feasibility Award (Su.K), NIH-NIDDK-R01DK080684 (S.S), the computational and data resources and staff expertise provided by Scientific Computing and Data at the Icahn School of Medicine at Mount Sinai (L.A.P), the Clinical and Translational Science Award (CTSA) grant UL1TR004419 from the National Center for Advancing Translational Sciences (L.A.P). The Advanced Light Microscopy Core (RRID:SCR_022526) and instruments used in this project were funded, in part, by the Pennsylvania State University College of Medicine via the Office of the Vice Dean of Research and Graduate Students and the Pennsylvania Department of Health using Tobacco Settlement Funds (CURE).

## AUTHOR CONTRIBUTIONS

M.B. conceived and led the project, designed experiments, analyzed data and wrote the manuscript; Sr.K., C.S., K.Sz. designed and executed experiments, analyzed data and contributed to writing the manuscript. A.Z.A, B.K., C.D. and L.N. performed experiments. J.P. and Su.K. contributed to the conceptual development of the project, and designed and oversaw scRNAseq experiments. M.S. and S.N. performed scRNAseq experiments. N.S. advised on bulk RNAseq experiments. C.B. advised on microscopy analysis. S.S. provided expertise in adult enteric neuronal culture. L.A.P. led the computational data generation and analysis and network analysis. Y.L. designed the pipeline and performed the analysis of bulk and single cell RNAseq data and generated the plots for the network analysis. W.W. and J.Z. constructed the Bayesian network models.

## COMPETING INTERESTS

The authors declare no competing interests.

## Notes

### Competing Interest Statement

The authors have declared no competing interest.

